# Inflammatory Mesenchymal Stromal Cells and IFN-responsive T cells are key mediators of human bone marrow niche remodeling in CHIP and MDS

**DOI:** 10.1101/2024.11.27.625734

**Authors:** Karin D. Prummel, Kevin Woods, Maksim Kholmatov, Eric C. Schmitt, Evi P. Vlachou, Gereon Poschmann, Kai Stühler, Rebekka Wehner, Marc Schmitz, Susann Winter, Uta Oelschlaegel, Logan S. Schwartz, Pedro L. Moura, Eva Hellström-Lindberg, Matthias Theobald, Jennifer J. Trowbridge, Uwe Platzbecker, Judith B. Zaugg, Borhane Guezguez

## Abstract

Somatic mutations in hematopoietic stem/progenitor cells (HSPCs) can lead to clonal hematopoiesis of indeterminate potential (CHIP), potentially progressing to myelodysplastic syndromes (MDS). Here, we investigated how CHIP and MDS remodel the human bone marrow (BM) niche relative to healthy elderly donors, using single cell and anatomical analyses in a large BM cohort. We found distinct inflammatory remodeling of the BM in CHIP and MDS. Furthermore, the stromal compartment progressively lost its HSPC-supportive adipogenic CXCL12-abundant reticular cells while an inflammatory mesenchymal stroma cell (iMSCs) population emerged in CHIP, which expanded in MDS. iMSCs exhibited distinct functional signatures in CHIP and MDS, retaining residual HSPC-support and angiogenic activity in MDS, corresponding with an increase in microvasculature in the MDS niche. Additionally, an IFN-responsive T cell population was linked to fueling inflammation in the stroma. Overall, these findings open new avenues for early intervention in hematological malignancies.

## Introduction

Aging is associated with a decline in hematopoietic system function and alterations in the composition of the bone marrow (BM) stem cell niche. Furthermore, it increases the risk for hematopoietic stem and progenitor cells (HSPCs) to accumulate somatic mutations in cancer-driver genes ^1^. Some of these mutations confer a competitive advantage and drive clonal expansion ^2^. This phenomenon, when the variant allele frequency (VAF) exceeds 2% in the absence of hematological neoplasms, is termed clonal hematopoiesis with indeterminate potential (CHIP) ^3,4^ and is detected in more than 10% of adults over 65 ^5,6^. Most recurrent CHIP mutations affect DNA methylation regulators (*DNMT3A, TET2, ASXL1*) and have been associated through a complex array of confounding factors with increased risks of non-malignant disorders such cardiovascular disease and rise in overall mortality ^7–10^. Importantly, CHIP may precede and increase the risk of developing hematologic malignancies such as myelodysplastic syndromes (MDS) and transformation to acute myeloid leukemia (AML) ^11–13^. As CHIP is increasingly recognized as a biomarker for the risk of blood malignancy development, understanding its role for transformation to MDS may provide a better tool for screening strategies and guiding clinicians in monitoring patients for signs of disease progression.

MDS represents a group of hematological malignancies arising from mutated HSPCs, resulting in ineffective hematopoiesis, BM dysplasia, peripheral blood cytopenia, and predisposition to secondary AML ^14^. While MDS shares some genetic features with CHIP, such as mutations in epigenetic regulators, it also exhibits cytogenetic aberrations and additional mutations, especially in RNA splicing factors (*SF3B1, SRSF2, ZRSR2*) ^15,16^. For instance, SF3B1 mutations are highly predictive for the occurrence of anemia and diagnosis of myeloid neoplasm ^17^. In particular SF3B1 mutations are found in 80% of low-risk MDS cases characterized with ring sideroblasts (RS), thereby defining a distinct MDS-SF3B1 subtype within MDS-RS ^18–20^.

The BM microenvironment (BM niche), including immune cells, endothelium and multipotent stromal cells (MSCs), constitutes specialized niches that orchestrate HSPC differentiation and maintenance, bone homeostasis, and immunity ^21,22^. These niches undergo remodeling with aging, marked by increased adipocytes (fatty BM) and a shift of MSC differentiation towards the adipogenic lineage ^23,24^; ^25^. Another hallmark of BM aging is the emergence of “inflammaging”, a sterile low-grade inflammatory state characterized by elevated levels of pro-inflammatory factors including IL-1β, IL-6, and TNFα ^26–28^. Inflammaging is associated with a decreased stem cell function ^29,30^ and has been observed across multiple organ systems ^31,32^. In CHIP carriers, elevated blood serum levels of IL-6, IL-8, IL-18, and TNFα have been detected compared to age-matched controls ^33–35^. Furthermore, the establishment of an inflammatory BM state has been shown to be beneficial for clonal expansion of *TET2*- and *DNMT3A*-mutant HSPCs in mouse models as well as in human CHIP cohort studies ^30,35–38^.

Extensive remodeling of the BM niche has been observed in hematopoietic malignancies, fostering an environment that supports malignant cells. This remodeling particularly influences the stromal components, which can alter the functions of resident immune cells, such as T cell-mediated responses ^39–41^. In MDS, *in vitro* cultured MDS-derived BM MSCs show an inflammatory program ^42^, decreased proliferation capacity, and reduced HSPC support ^43^ compared to healthy donor-derived BM MSCs. A recent study in a *DNMT3A*-mutant CHIP mouse model (equivalent to a human VAF of 1-4%) suggest that mutated HSPCs can induce senescence in BM MSCs through pro-inflammatory cytokines including IL-6 and TNFα ^44^. However, *in vivo* studies on the MDS BM niche cells remain scarce, and the extent to which CHIP-mutant clones can remodel the BM microenvironment remains unclear, particularly in the human context and at single-cell level.

In this study, we aim to elucidate the alterations in the BM microenvironment associated with CHIP and low-risk MDS. We assembled a cohort of aged control donors, CHIP carriers (DNMT3A, TET2), and low-risk MDS patients with overlapping mutations (TET2, DNMT3A, and spliceosome mutations). Using multimodal analyses, including targeted RNA profiling, flow cytometry, scRNA-seq, and immunofluorescent imaging of BM aspirates and matching FFPE bone biopsies, we delineated the cellular composition and functional states of the microenvironment to identify potential molecular drivers of disease progression. Furthermore, we applied *in vitro* co-culture systems to explore the stromal interactions with MDS cells, providing insights into the microenvironmental changes that may promote or sustain malignant hematopoiesis. Overall, our results highlight the emergence of an inflammatory MSC subset in CHIP that expands in MDS and plays alongside IFN-responsive T cells, a key role in BM niche remodeling and potentially contributing to disease progression.

Our findings contribute to a deeper understanding of the inflammatory human BM microenvironment in context of CHIP and MDS, highlighting the role of stromal and immune cell interactions with mutated HSPCs as key mediators in fueling disease-associated BM inflammation.

## Results

### Distinct inflammatory and immune microenvironment profiles in CHIP and MDS bone marrow

To understand how the microenvironment is altered across CHIP and MDS with overlapping somatic mutations in the human context, we analyzed bone marrow (BM) samples from a balanced cohort of 84 donors, including 35 age-matched controls, 17 CHIP donors (VAF ≥ 2%), and 32 low-risk MDS patients from newly diagnosed or disease-modifying treatment-naïve cases (i.e. erythropoietin or supportive care only) (**Fig. 1A**). Many CHIP and MDS donors carried the most prevalent *DNMT3A* and *TET2* mutations. In addition, the majority of MDS patients also harbored at least one mutation in a spliceosomal gene, such as *SF3B1* (16 patients), *SRSF2*, or *U2AF1* (6 patients) or carry only *DNMT3A/TET2* mutations (10 patients) (**Supplementary Table 1, Fig. S1A**). Blood count assessment showed equal proportions of leukocytes across the mutational spectrum of the MDS cohort but indicated the presence of increased ring sideroblasts (RS) and signs of anemia only in MDS carrying *SF3B1* mutations (**Fig. S1B**), which is consistent with the current WHO consensus classification of MDS-RS ^45,46^.

**Figure 1:**
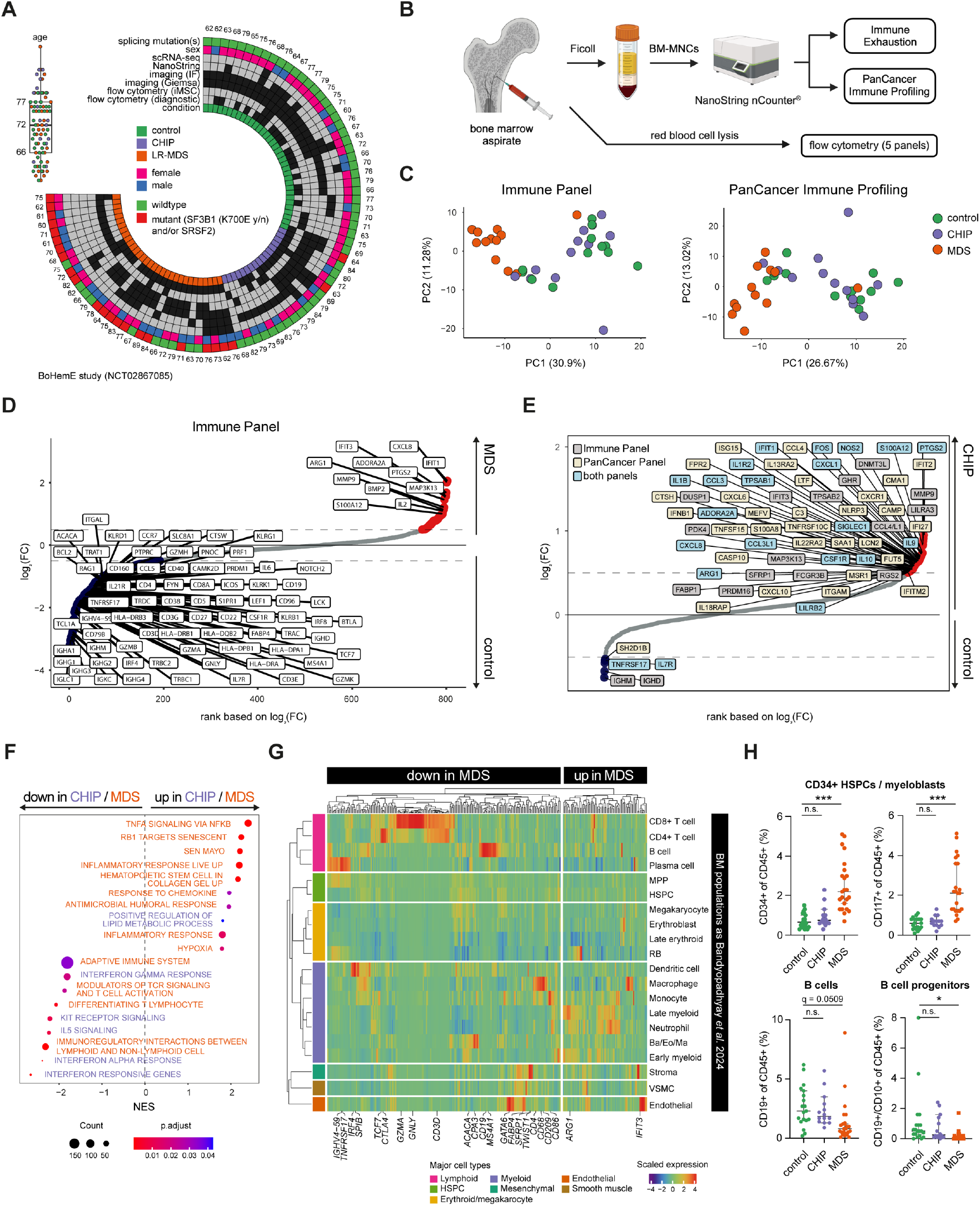
Expression profiling in bulk bone marrow reveals progressive inflammation from CHIP to MDS. **(A)** Circular heat map displaying the overview of our study using the BoHemE cohort (NCT02867085), including age distribution (boxplot on the top left) and other attributes such as sex, splicing mutations (*SF3B1, SRSF2*), and conditions (green: control, purple: CHIP, orange: MDS). The rings indicate inclusion (black boxes) in various analyses such as scRNA-seq, NanoString, flow cytometry, and immunofluorescent imaging. **(B)** Schematic summarizing the experimental workflow. Bone marrow (BM) aspirates were processed into BM mononucleocytes (BM-MNCs) using Ficoll separation or only red blood cells were lysed. Ficoll-treated samples were used for NanoString immune profiling (Immune Exhaustion and PanCancer panels) and red blood cell lysed samples were used for flow cytometry analysis to quantify major cell types (see **Supplementary Table 2** for NanoString panels and **Supplementary Table 3** for flow cytometry panels). **(C)** Principal component analysis (PCA) scatter plots showing the separation of control, CHIP, and MDS samples based on the Immune panel (left) and PanCancer Immune panel (right). Each dot represents a sample, with colors indicating condition (green: control, purple: CHIP, orange: MDS). **(D-E)** Rank plot showing the log2 fold changes in gene expression between MDS and control samples (D) and CHIP versus control samples (E). Genes with an absolute log2 fold change > 0.5 are labeled. Significant genes are highlighted in red and in E) with color-coded boxes around the gene names indicating the NanoString panel (Immune or PanCancer). **(F)** Gene Set Enrichment Analysis (GSEA) results illustrating gene sets significantly upregulated or downregulated in MDS versus control (orange) and CHIP versus control (purple). **(G)** Heatmap displaying the expression levels of NanoString-derived genes that are differentially expressed between MDS and control samples in D) on scRNA-seq data from healthy BM ^47^. Genes are averaged by cell type and scaled across the cell types for visualization. **(H)** Flow cytometry analysis comparing the relative proportions of HSPCs/myeloblasts characterized by CD34 or CD117, CD19^+^ B cells, and CD10^+^CD19^+^ B cell progenitors within the CD45^+^ cell population across control (green), CHIP (purple), and MDS (orange) donors. Median with 95% confidence intervals is shown. Statistical significance: n.s. = not significant; * q < 0.05; ** q < 0.01; *** q < 0.001 (one-way ANOVA, FDR for multiple comparison correction).

To gain a comprehensive overview of immune microenvironmental alterations, we profiled gene expression on total MNCs from bulk BM aspirates using two NanoString nCounter panels targeting 773 immune-related and 730 cancer inflammation-associated genes (**Supplementary Table 2, Fig. 1B**). Principal component analysis (PCA) of these data revealed a separation of MDS samples from CHIP and controls along the main principal component for both panels (**Fig. 1C; Methods**), indicating a global remodeling of the BM immune milieu in MDS. CHIP did not differ substantially from control along the main principal components, but within the two groups, 8 CHIP/control donors clustered separately (**Fig. 1C**). Differential gene expression analysis identified 123 unique genes (FDR < 0.05) between MDS and controls, with the majority being downregulated in MDS BM and only 17 genes upregulated (**Fig. S2A**,**B**). Notable upregulated genes in MDS included inflammation-associated genes, including the interferon-induced family of genes *IFIT1/2/3*, while downregulated genes were involved in antigen-presentation (e.g., IGH^-^ and HLA-associated genes), antigen-recognition (*CD3*), and B cell lineage markers (*CD38, CD19*; **Fig. 1D**). GO enrichment analysis of downregulated genes in MDS relative to the controls further highlighted a disturbed regulation and activation ability of T cells in MDS (**Fig. S2C**). Gene set enrichment analysis (GSEA) revealed upregulation of TNFα and IFNα pathways in MDS compared to control and CHIP, indicating a pro-inflammatory environment (**Fig. S2D**,**E**). Although no statistically significant changes were found between CHIP and controls, a ranking of log2 fold changes (log2FC) without an FDR cutoff highlighted moderate upregulation of 63 genes with a log2FC of 0.5, including *PTGS2, IL1B, IL1R1/2, CXCL8*, and *TNF* in CHIP (**Fig. 1E**). In contrast to MDS, B- and T cell lineage markers were not downregulated in CHIP (**Fig. 1E**). Based on clonal size (VAF), CHIP samples with VAF ≥ 5% formed a distinct cluster that included mostly DNMT3A-mutated cases, along with a few control donors (**Fig. S2F**). This cluster showed increased expression of inflammatory and proliferative markers compared to the other CHIP/control cluster (VAF < 5%), rendering it more similar to the MDS samples (MDS-like, **Fig. S2G)**.

To identify the cell types contributing to the global changes in the MDS niche, we assessed the cell type-specific expression of the differentially expressed genes from the NanoString analysis in a recently published single-cell atlas of healthy BM ^47^. Specifically, for each differentially expressed gene we averaged the expression in each healthy BM cell type and clustered the resulting gene-by-cell expression matrix (**Fig. S2H**). Based on this, we grouped cell types exhibiting similar patterns into the major BM cell populations and reclustered the genes (**Fig. 1G**). This revealed that genes downregulated in MDS were predominantly expressed in T cells, pre-B and pro-B cells, B cells, and B plasma cells (e.g., *CD3, CD8, CD4, CD22, IGH, CD38, CD19*) (**Fig. 1G**).

To determine whether the reduced gene expression reflected a lower abundance of these cell populations within the MDS BM, we compared the transcriptional profiling with immunophenotyping data of the major BM cell populations at the time of diagnosis (**Fig. 1H, Fig. S1C**). This confirmed a reduction in B cells in MDS patients compared to controls, consistent with prior studies on low-risk and SF3B1-mutated MDS ^48–51^, while no significant changes were observed in other cell populations, including T cells, granulocytes, NK, and NKT (**Fig. S1C, Supplementary Table 3, Supplementary Table 4**). Despite flow cytometry showing no change in overall T cell numbers **(Fig. S1C)**, several pro-inflammatory genes upregulated in MDS BM (e.g., *TNF, IFNL1, CCR9*) were predominantly expressed in T cells (**Fig. 1G**), suggesting the presence of a distinct pro-inflammatory T cell population in LR-MDS. No significant differences between control and CHIP were observed (**Fig. 1G, S1C**). Flow cytometry further confirmed an increase in CD34+/CD117+ HSPCs, likely driven by the presence of immature myeloblasts in MDS (**Fig. 1H**).

Additionally, the lower expression of genes specific to dendritic cells in MDS samples, and higher expression of genes specific to neutrophils and myeloid progenitors suggests the functionality or abundance of these populations was also affected (**Fig. 1G**). The gene expression in the stromal and endothelial compartments also showed distinct patterns, with some stromal genes (e.g., *TWIST, SFRP1*) being more highly expressed in controls, while other (e.g., *BMP2*) were upregulated in MDS (**Fig. 1G**). In endothelial cells, genes such as *IFIT3* were elevated in MDS (**Fig. 1G**). Since these genes are reported as important modulators in HSPC and osteogenic differentiation ^52–55^, our data indicated that the stromal and endothelial compartments might also undergo significant remodeling in the BM of MDS patients.

Overall, the bulk expression analysis of BM suggests an increasingly pro-inflammatory microenvironment from CHIP to MDS, characterized by the expansion of immature myeloid cells, loss of B cells, and selective activation of inflammatory pathways, particularly involving TNFα and IFNα signaling.

### Single cell RNA-seq reveals inflammatory stromal and lymphocyte subsets in the BM niche of CHIP and MDS

Building upon the immune and inflammatory gene expression changes observed in the targeted transcriptome analysis of whole BM cells **(Fig. 1)**, we employed single cell transcriptomics to profile key cell types suspected to contribute to the BM niche remodeling in CHIP and MDS. We focused on stromal cells, T cells, as well as (mutated) HSPCs from a representative subset of the cohort, which included 3 controls, 3 CHIP (DNMT3A and/or TET2), and 4 MDS patients (SF3B1/SRSF2 and DNMT3A and/or TET2) (**Fig. 2A, Fig. S1A**) more info in **Supplementary Table 5**).

**Figure 2:**
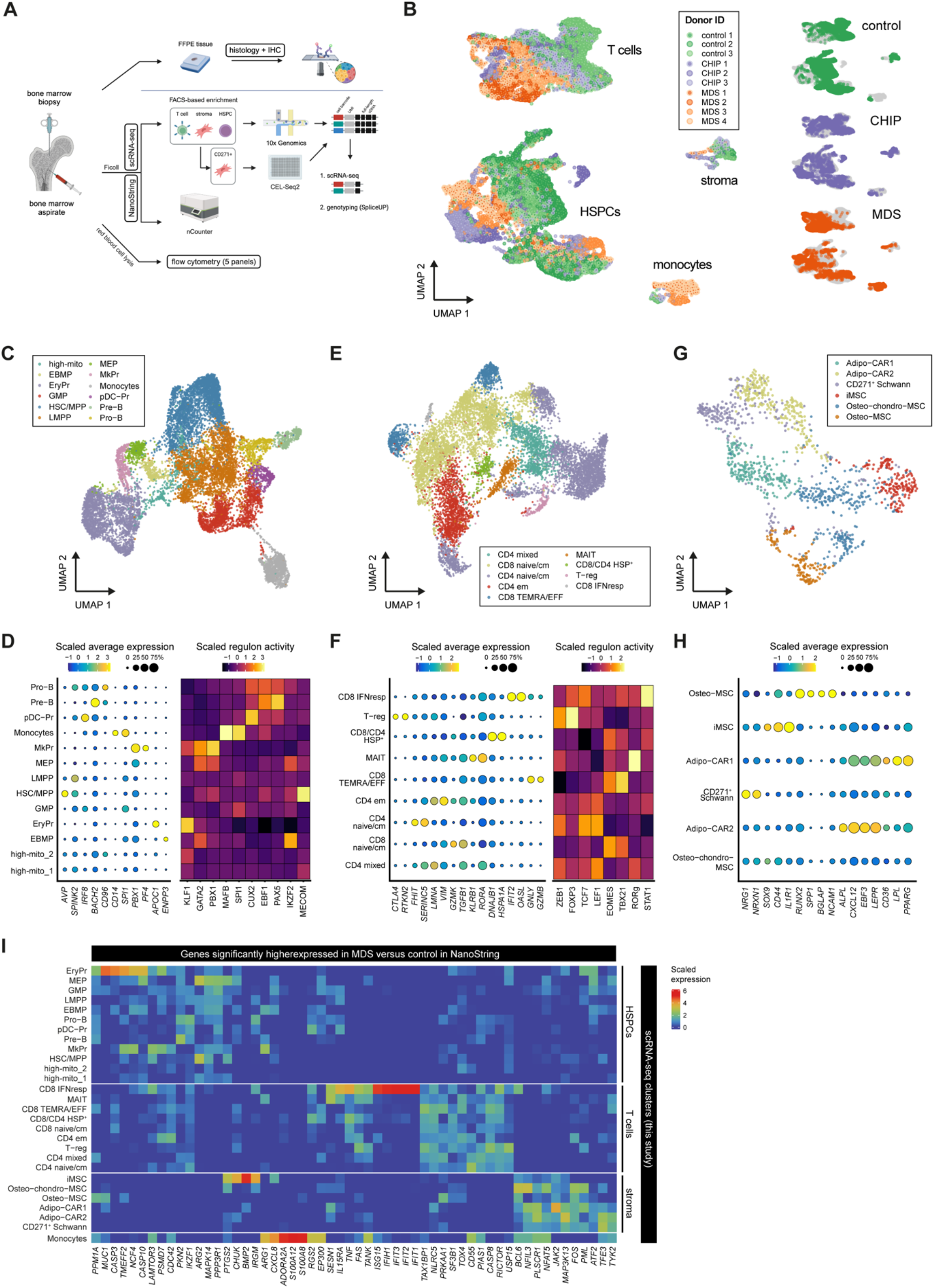
Identification of cell type subsets of BM in healthy, CHIP, and MDS cohort by scRNA-seq. **(A)** Schematic of the experimental design performed with the donor-derived BM aspirates and FFPE trephine bone biopsies. **(B)** Uniform manifold approximation and projection (UMAP) of all major cell populations of scRNA-sequencing of donors highlighted in **Fig. 1A** and **Fig. 2A**. Cells are colored by Donor ID and split by condition. **(C)** UMAP of HSPCs, with clusters defined and colored by cell type annotation. **(D)** Expression of marker genes of individual cell types within the HSPC population. Bubbles in dot plot correspond to direct expression of marker genes with sizes corresponding to percent of cells expressing the given gene. The heatmap shows average AUC (area under the curve) scores of marker transcription factors (TFs) based on regulons inferred in the current study. **(E)** UMAP of T cells, with clusters defined and colored by cell type annotation. **(F)** Expression of marker genes of individual cell types within the HSPC population. Bubbles correspond to direct expression of marker genes with sizes corresponding to percent of cells expressing a given gene. The heatmap shows average AUC scores of marker TFs based on regulons from ^65^. **(G)** UMAP of stromal cells integrated between 10x data and CellSeq2 colored by cell type annotation. **(H)** Markers of individual cell types within the stromal population. **(I)** Average expression levels of genes that were significantly higher expressed in MDS compared to control in the NanoString data, across the cell type clusters found in our scRNA-seq dataset.

To achieve a comprehensive representation of the cell types of interest without lineage over-representation (such as immature cells from MDS-RS), we used FACS to enrich for 3 fractions: non-hematopoietic stromal fraction (using a negative gating strategy: CD45^-^CD235a^-^ CD71^-^CD14^-^CD38^-^), HSPCs (CD45^+^CD34^+^CD235a^-^CD71^-^CD14^-^), and T cells (CD45^+^CD3^+^CD235a^-^CD71^-^CD14^-^) (**Fig. S3A**).

The HSPC and T cell fractions were combined and subjected to scRNA-seq (10x Genomics) in parallel with the stromal fraction, ensuring clean separation of the populations and minimizing contamination of blood or immune cells in the stromal fractions. To enhance stromal cell recovery, we additionally sorted CD271^+^ cells, an established marker of human stromal cells ^56,57^ that was marking the biggest cell population compared to other described human BM stromal markers (**Fig. S3B**,**C**), for plate-based scRNA-seq (CEL-Seq2).

After donor demultiplexing based on genotype deconvolution (**Methods**), quality control, and cell calling (**Fig. S3D-H**), we identified a population of hematopoietic cells identified in the stromal gate despite the removal of CD38^+^ and CD71^+^ cells (**Fig. S3I; Methods**). These cells resembled monoblasts and erythroblasts (or RS) and predominantly originated from MDS donors and were removed for the main analyses. Altogether, these filtering steps yielded 24,693 cells, with between 1,190 and 4,393 cells per donor (**Supplementary Table 4**). A UMAP representation followed by broad cell type classification revealed clear clustering of the cell populations of interest: 1,396 stromal cells (*CXCL12* and *LEPR*), 9,962 T cells (*CD3E* and *CD247*), and 12,257 HSPC (*CD34* and *SPINK2*) from all donors (**Fig. 2B**). Notably, we also recovered a population of CD14-dim monocytes (low *CD14* and *LYZ*), which was unexpected given our stringent sorting strategy for CD14^-^ cells. These monocytes primarily originated from MDS samples (MDS1 and MDS3). After filtering, we retained cells with high-quality metrics (median 3,644 UMIs, 1,610 unique genes, and 6.2% mitochondrial reads per cell), ensuring that all major cell types were well-represented across all conditions (**Supplementary Table 5**).

To further refine the cell populations, we integrated the scRNA-seq data across all donors and for each major cell population separately using Seurat v3 (see **Methods**) and generated a UMAP embedding for each broad cell type (T cells, stroma, HSPCs including CD14-low monocytes). We next performed Leiden clustering and annotated the populations by manually curating known marker genes and referencing well-annotated human T cells and HSPC atlases (see **Methods** ^58–65^). To refine the annotation for some cell types, we also inferred their main transcription factor regulons using SCENIC ^66^ and cross-referenced the results with publicly available markers (**Methods**, ^60–63,66^). In total, we defined 13 HSPC clusters, 9 T cell clusters, and 6 stromal cell clusters (**Fig. 2C-H**). For the HSPC subsets we identified HSC and multipotent progenitors (HSC/MPP, *HLF, AVP, GPR56, MECOM*), lymphoid-primed multipotent progenitors (LMPP, *SPINK2, CD34, PROM1*), B-lymphoid progenitors (Pro-B, *CD96, DNTT, EBF1* and Pre-B cells *BACH2, PAX5*), myeloid progenitors (GMP, *PRTN3, MPO* and pDC-Pr, *IRF8, CUX2*), and megakaryocyte-erythroid progenitors (EBMP, *ENPP3, IKZF2;* MEP, *GATA2;* MkPr, *PF4, PBX1* and EryPr, *APOC1, KLF1*). A myeloid population reminiscent of non-classical CD14-dim monocytes (*CD68, MFAB, SPI1*) was also identified. These were previously defined as patrolling monocytes involved in the innate local surveillance of tissues and shown to be increased in blood malignancies ^67–69^. Besides that, two mixed clusters were detected that contained relatively high levels of mitochondrial genes (high_mit) or were specific to an individual donor (Control 3) (**Fig. 2C**,**D**).

We annotated 9 T cell populations based on previous studies using a combination of marker genes and regulon activities of specific transcription factors, including those inferred from SCENIC and from our previously established T cell-specific regulons ^63,65,70,71^. The CD4 clusters included: CD4 naive/central memory (CD4_naive/CM_, *SELL, CCR7*), CD4 effector memory (CD4_EM_, *IL7R, KLRB1, AQP3*), a mixed CD4 population, and regulatory T cells (Treg, *IL2RA, RTKN2*, FOXP3 regulon activity). Among CD8 T cells, we distinguished CD8 naive/central memory (CD8_naive/CM_, *CCL5, GZMK*), CD8 terminally differentiated effector memory re-expressing *CD45RA* (T_EMRA_, *FCGR3A, GZMB, GNLY*), as well as IFN-responsive CD8 effector cells (CD8 IFN response, *OASL, IFIT2*). Additionally, we observed a mixed CD4/CD8 heat-shock protein-expressing population (CD8/CD4 HSP^+^, *DNAJB1, HSP90AA1, HSPA1A*) and mucosal-associated invariant T cells (MAIT, *SLC4A10, ME1*) (**Fig. 2E**,**F)**.

Within the stromal compartment, we annotated 6 distinct subpopulations based on BM reference atlas ^47^, including two adipogenic CXCL12 abundant reticular (adipo-CAR1/2, *CXCL12, LEPR, EBF3, PPARG*), chondrogenic-(*SOX9*), and osteogenic-lineage (*RUNX2, SPP1, NCAM1, BGLAP)* MSCs, as well as neural crest-derived Schwann-like cells (*NRG1, NRXN1*), which express *CD271 (NGFR)* akin to MSCs ^72,73^ and were mostly identified among our CD271-enriched population (**Fig. 2G**,**H**). The two Adipo-CAR populations can be distinguished by distinct adipose-lineage markers: Adipo-CAR1 (*PPARG, LPL*) and Adipo-CAR2 (*ALPL, PDGFRB*), reminiscent of the *Lepr*-expressing Adipo-CAR populations identified in mouse BM ^74,75^. Of particular interest was a population expressing inflammation-associated genes, including *CD44* and *IL1R1* ^76,77^. We refer to these stromal cells as inflammatory MSCs (iMSCs). Other stromal cells, including endothelial cells, vascular smooth muscle cells / pericytes, and mature osteoblasts and adipocytes could not be recovered from BM aspirates, either due to their low abundance or mechanical fragility during cell isolation.

Integrating the scRNA-seq with our NanoString expression data from the larger cohort revealed MDS-specific cell populations that contribute to the elevated inflammatory gene expression in MDS, which were absent in the healthy BM reference atlas (**Fig. 2I**). Notably, genes expressed in iMSCs, IFN-responsive T-cells, and the non-classical CD14-low monocyte population were upregulated in MDS BM, suggesting the presence of cell types or cell states in the MDS-remodeled BM niche that are not apparent in the healthy bone marrow.

Overall, our single cell data reveals the cellular variety of HSPC, stromal, and T cell compartments across the CHIP and MDS BM. Integration with the bulk transcriptomics data of the larger cohort highlights the emergence of inflammatory populations in MDS, particularly iMSCs and IFN-responsive T cells, underscoring their potential role in BM niche remodeling.

### Inflammatory MSCs are exclusively present in CHIP and MDS

Next, we focused specifically on the stromal cell populations and quantified the changes in cell subtypes across control, CHIP, and MDS BM among the enriched stroma. The most notable shift was the emergence of the iMSC population, exclusively present in CHIP and MDS (**Fig. 3A, Supplementary Table 6**). This iMSC population was detected in 2 out of 3 CHIP donors and all MDS donors examined, albeit in varying proportions (**Supplementary Table 5**). Despite the absence of a significant pro-inflammatory signature in whole BM from CHIP donors (**Fig. 1E**,**F**), our scRNA-seq workflow successfully identified these very rare iMSCs in CHIP, highlighting the sensitivity of our approach.

**Figure 3:**
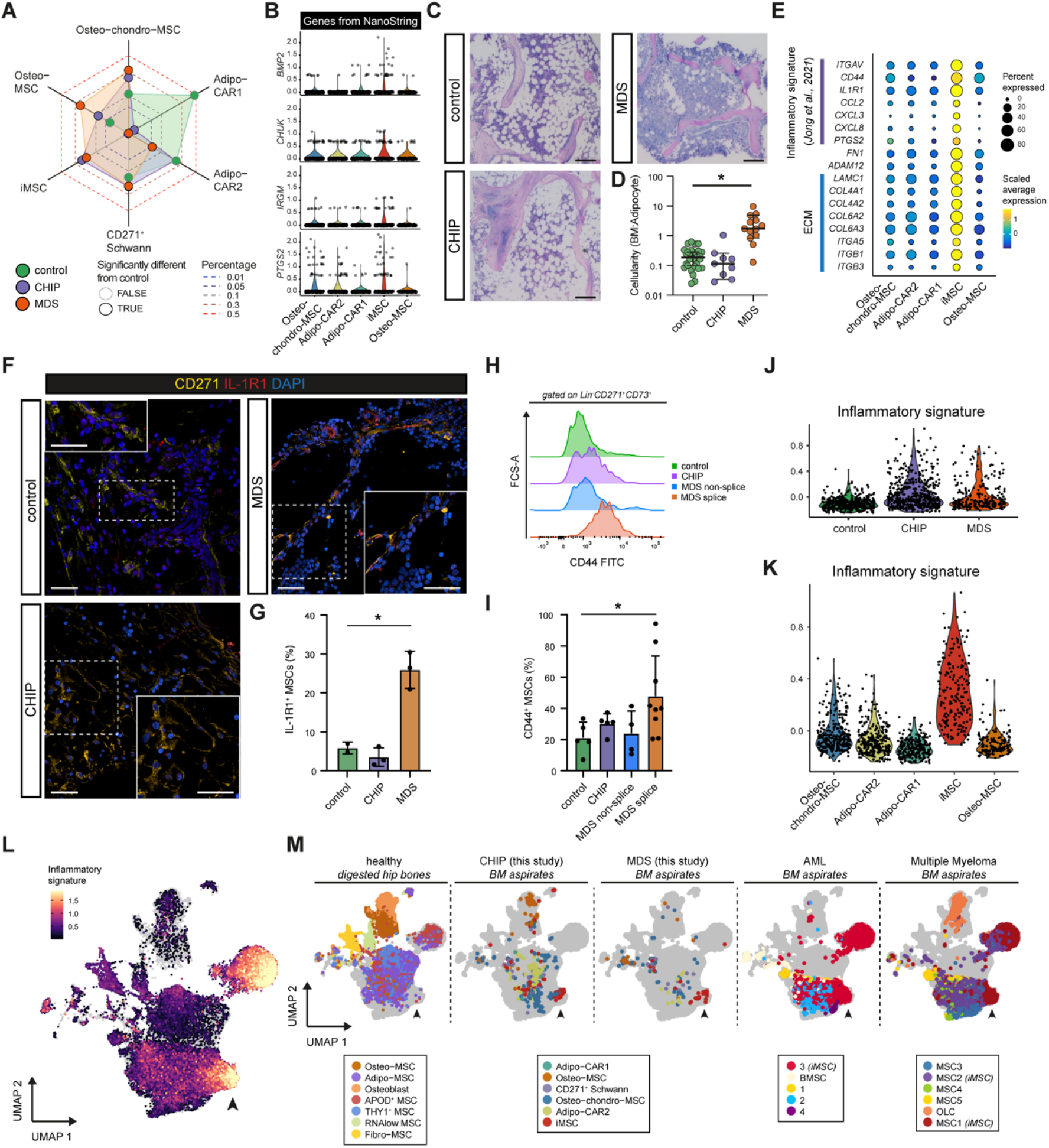
Identification of a distinct inflammatory MSC population in BM of CHIP and MDS. **(A)** Spider plot displaying the relative abundance of cell types found in the stromal population between different conditions control (green), CHIP (purple), and MDS (orange) donors. Cell type proportions are plotted to the radius after probit transformation, with a black outline on points indicating statistically significant differences compared to controls (see **Methods**). **(B)** Violin plots show the expression levels of key stromal genes that are significantly upregulated in MDS in the NanoString across the stromal subtypes. **(C)** Representative Giemsa staining images of bone marrow sections from control, CHIP, and MDS patients. Scale bars are 100 μm. **(D)** Quantification of cellularity (see also **Fig. S4A-C**), a ratio of the BM cells and adipocytes, in each condition is shown as a scatter plot, with median values indicated. Statistical significance: * p < 0.0001, Kruskal-Wallis test. **(E)** A dot plot showing the expression levels of selected ECM-related and inflammatory markers (from ^76^) across different stromal cell populations in the bone marrow, specifically highlighting the iMSCs. Circle size represents the percentage of cells expressing the gene, while color intensity represents scaled expression. **(F)** Immunofluorescence images show IL-1R1 (red) and CD271 (yellow) co-staining in BM FFPE tissue sections from control, CHIP, and MDS patients. Insets highlight areas containing inflammatory IL-1R1^+^CD271^+^ stromal cells. DAPI (blue) stains the nuclei. Scale bars are 25 μm. **(G)** Bar graph quantifying the percentage of IL-1R1^+^ MSCs across control, CHIP, and MDS samples (n = 3 / condition). MDS shows a significant increase in IL-1R1^+^ MSCs. Statistical significance: * q < 0.05, one-way ANOVA (FDR for multiple comparison correction). **(H)** Histogram showing the distribution of CD44 expression on Lin^−^ CD271^+^CD73^+^ MSCs from control, CHIP, and MDS samples as measured by flow cytometry. MDS samples are divided into splice and non-spliceosome (*SF3B1, SRSF2*) mutated categories for further analysis. **(I)** Bar graph quantifying the percentage of CD44^+^ MSCs in control, CHIP, and MDS samples from H). Mean with SD is shown. Statistical significance: * q < 0.05 (one-way ANOVA, FDR for multiple comparison correction) **(J-K)** Violin plots showing the distribution of inflammatory signature scores in stromal cells found in our scRNA-seq dataset (I) across conditions (control, CHIP, MDS) (J) and across distinct stromal cell subtypes. The signature score represents the cumulative expression of inflammation-related genes. **(L)** UMAP projection is showing the integration of our scRNA-seq dataset with 3 published human BM datasets of healthy BM, Acute Myeloid Leukemia (AML) BM, and Multiple Myeloma (MM) BM ^47,76,85^ using Harmony ^200^. The stroma cells are colored by their inflammatory signature scores. 51.95% of the cells expressed genes (part) of the inflammatory signature. Higher signature scores (yellow) are associated with inflammatory stromal subtypes. Arrowhead marks iMSCs. **(M)** Comparative UMAP plots of stromal cell populations from healthy, CHIP, and MDS BM aspirates (left), healthy digested hip bones (middle) ^47^, and bone marrow aspirates from AML ^85^ and MM patients (right) ^76^. Cluster annotations are taken from their respective original publications, but the inflammation-associated clusters are annotated with “iMSC”. Arrowheads mark iMSCs.

Notably, the iMSCs expressed several genes significantly upregulated in the NanoString transcriptomics of bulk MDS BM (**Fig. 3B**), suggesting that even bulk expression analysis picked up inflammatory signatures from this population in BM aspirates. Key genes expressed include *CHUK* (encoding IKK-alpha, crucial for NF-κB signaling activation and regulation), *BMP2*, and *PTGS2* (a prostaglandin synthase involved in the inflammatory response) ^78,79^.

In addition to the expansion of iMSCs, MDS patients exhibited a loss of both Adipo-CAR1 and Adipo-CAR2 MSCs, while CHIP donors specifically lost the Adipo-CAR1 population. This absence of adipo-MSCs in MDS aligns with the downregulation of adipocyte-specific markers (*ACACA, FABP4*) in MDS in the NanoString data (**Fig. 1I**), suggesting an overall loss of adipocytes in MDS BM. This was confirmed by BM imaging, showing a reduction of BM adipocytes in MDS (**Fig. 3C**,**D, Fig. S4A-C, Supplementary Table 7**), which is in accordance with previous histological reports linking this shifted ratio to the expansion of the ineffective erythroid/myeloid compartment in LR-MDS ^80–82^.

Given the striking difference in the abundance of stromal populations in CHIP and MDS compared to controls, we further characterized the CHIP- and MDS-enriched iMSC population. These iMSCs displayed strong expression of inflammatory genes (*CD51, CD44, IL1R1, PTGS2, CXCL8, CCL2*) and pro-inflammatory extracellular matrix (ECM) remodeling factors (*ADAM12, FN1, LAMC1, ITGA5, ITGB1, ITGB3*) (**Fig. 3E**). They also expressed profibrotic canonical collagen IV and VI genes (*COL4A1, COL4A2, COL6A2, COL6A*3). The presence of iMSCs was further confirmed *in situ* by immunofluorescent staining of IL1-R1 in FFPE BM tissue sections, which revealed higher IL1-R1^+^ cell counts in MDS BM compared to control and CHIP donors (**Fig. 3F**,**G**). Despite the overall low IL1-R1^+^ cell count in CHIP, we could identify rare clusters of IL1-R1^+^/CD271^+^ cells in CHIP FFPE BM tissues (**Fig. S5A**). Additionally, flow cytometry with CD44 and CD51/61 on a subset of the control, CHIP, and MDS donors (**Fig. 1A**) further validated the appearance of iMSCs: We observed a significant increase of CD44^+^ expressing CD271^+^/CD73^+^ stromal cells in MDS BM aspirates, with this increase predominantly seen in spliceosome factor-mutated MDS (**Fig. 3H**,**I, Fig. S5B-D**). CD44 is linked to stromal inflammation and ECM production; its activation stimulates multiple downstream pathways including TGFβ and RhoA-YAP signaling pathway ^83,84^.

Inflammatory alterations in stromal cells have been previously reported in other myeloid malignancies such as AML ^85^ and multiple myeloma ^76^, linking these changes to cancer-associated fibroblast signatures ^86^. An inflammatory signature based on these previous studies (21 genes, **Supplementary Table 8, Methods**) confirmed that in our study inflammation was absent in MSCs from controls (**Fig. 3J**) and that the CHIP/MDS-specific iMSCs showed the highest expression-based scores for the inflammatory MSC signature (**Fig. 3K**). This was confirmed by integrating our data with existing healthy BM, multiple myeloma, and (NPM1-mutant) AML patients stromal data sets (**Methods**, ^47,76,85^, which revealed a distinct iMSC cluster shared between CHIP and MDS, and representing a subset of the iMSCs found in AML and Multiple Myeloma (MM) (**Fig. 3L**,**M, Fig. S6A**). This shared iMSC cluster across multiple myeloid malignancies exhibited a high inflammatory signature (**Fig. 3L, Fig. S6A**). Notably, iMSCs were absent except for a few cells in the healthy BM ^47^ and controls from the MM and AML studies ^76,85^, suggesting a shared stromal response to (chronic) inflammatory stimuli in myeloid diseases.

These findings suggest a slightly increased inflammatory niche in CHIP and a more pronounced expansion of an iMSC subset in MDS and AML.

### iMSCs differ between CHIP and MDS in their HSPC-support signatures

Next, we aimed to understand how HSPC support is affected in the remodeled and inflammatory niche. To investigate this, we generated an HSPC-support factor gene signature (46 genes, **Supplementary Table 5**) based on the literature ^87^ and assessed its expression across the different stromal populations. We observed that the adipo-MSC populations (Adipo-CAR1+2) showed the highest HSPC support signatures (**Fig. 4A**). Given the loss of these populations in MDS, and to a lesser extent in CHIP (**Fig. 3A, Fig. S6B**), this indicates a reduced capacity of the stromal compartment to maintain HSPCs in CHIP, which is even further pronounced in MDS. Indeed, comparing the *in situ* expression of CXCL12, a HSPC support and maintenance factor ^88^, relative to the number of CD271-expressing stromal cells by immunofluorescence on a selection of trephine BM biopsies from our CHIP/MDS cohort, we observed a trend towards lower CXCL12/CD271 ratio in MDS compared to control and CHIP (**Fig. 4B**,**C**). This suggests that either MSCs express lower levels of CXCL12 or the proportion of CXCL12-expressing MSCs within the CD271-positive MSC population was smaller in MDS. This is consistent with the overall decrease of HSPC-support signature expression in the scRNA-seq in CHIP and MDS when considering all stromal cells (**Fig. 4D**).

**Figure 4:**
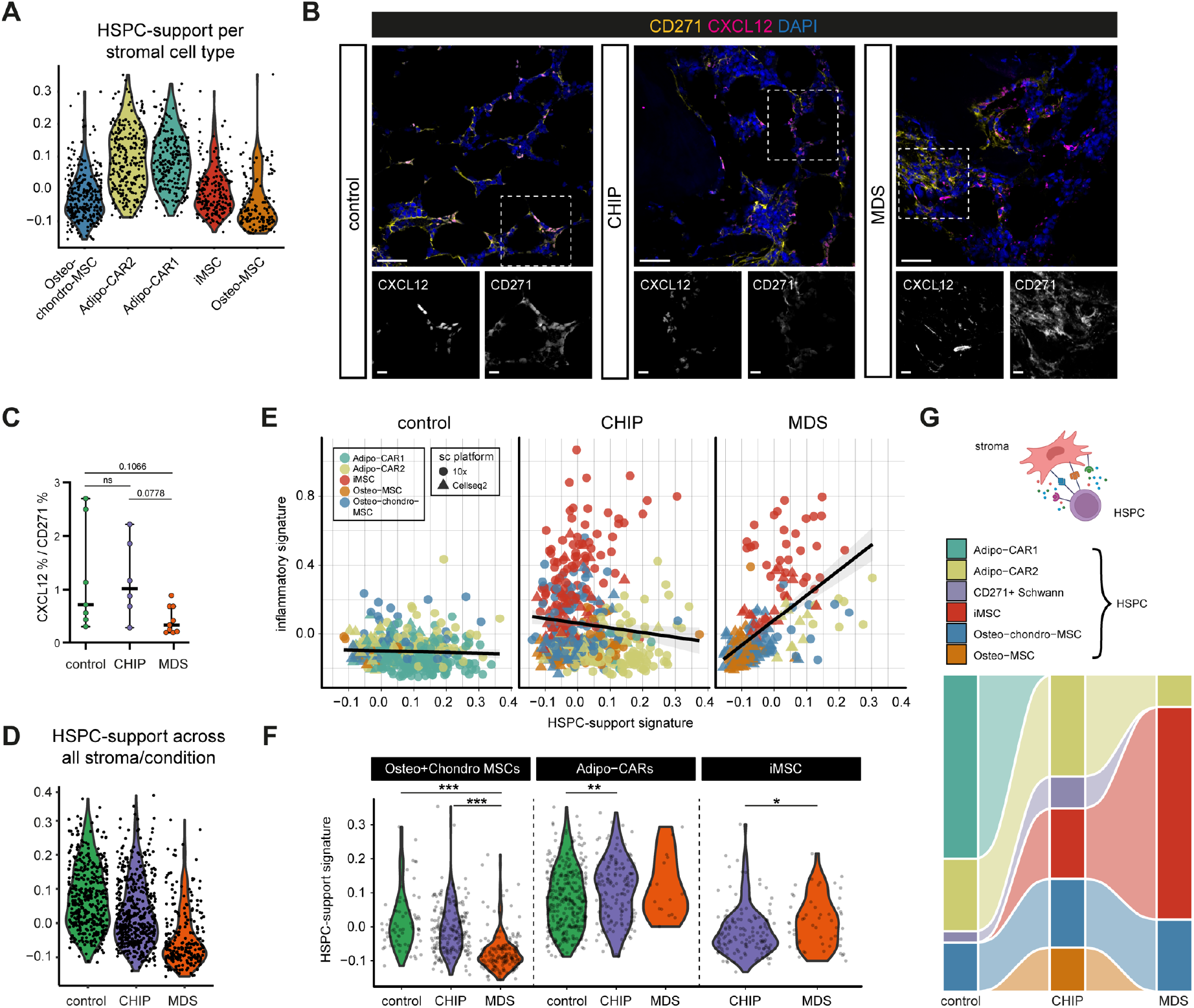
Mapping HSPC-support in stromal BM populations. **(A)** Violin plots showing the distribution of HSPC-support signature scores across the stromal cell types. **(B)** Representative immunofluorescence images showing CXCL12 (magenta) and CD271 (yellow) expression in control, CHIP, and MDS FFPE BM tissue sections. Insets highlight key regions of interest with distinct co-localization patterns. DAPI (blue) stains the nuclei. Scale bars are 25 μm. **(C)** Strip plot showing the percentage of CXCL12^+^ cells among CD271^+^ stromal cells. Middle bars represent the median. Statistical test: ns = not significant, adjusted P-values, Kruskal-Wallis test. **(D)** Violin plots showing the distribution of HSPC-support signature scores in stromal cells found in our scRNA-seq dataset across conditions (control, CHIP, MDS). The signature score represents the cumulative expression of the HSPC-support relative to controls. **(E)** Scatter plots showing the relationship between HSPC-support signature and inflammation signature scores across stromal cells between conditions. Different colors represent the distinct stromal cell populations. Black trend lines indicate the direction of the correlation in each condition. **(F)** Violin plots comparing the HSPC-support signature scores specifically in Osteo-chondro MSCs and Adipo-CAR1/2 across all conditions, as well as in iMSCs between CHIP and MDS. Statistical significance: * p < 0.05, ** p < 0.01, ** p < 0.005, Wilcoxon rank-sum test). **(G)** Sankey diagram summarizing the proportions of inferred cell-cell interactions inferred by NICHES ^89^ between HSPCs and different stromal cell types across control, CHIP, and MDS samples. Interaction proportions highlight the loss of supportive interactions in MDS.

Notably, in CHIP-derived stromal cells, we observed a slight anti-correlation between the HSPC-support and the inflammatory signatures (**Fig. 4E**), suggesting that iMSCs in CHIP can provide little HSPC support. In contrast, in MDS, the iMSCs retained residual expression of hematopoietic support genes, resulting in a positive association between the inflammatory and HSPC support signatures across all MSC populations (**Fig. 4E**). Indeed, we observed a higher support signature in iMSCs from MDS, including expression of *CXCL12* and *KITLG* (**Fig. S6C**), compared to iMSCs from CHIP, although the overall support provided by the iMSCs was still lower than in the Adipo-CAR1/2 populations in controls and CHIP (**Fig. 4F**). The same analysis on the AML stroma from Chen and colleagues ^85^ revealed a similar trend as seen in MDS (**Fig. S6D**). GSEA of iMSCs from MDS versus CHIP showed an upregulation of pathways related to UV response, apoptosis, mTORC1 signaling, and notably TNFα signaling via NF-κB (**Fig. S6E**), highlighting the progressive increase of stress and inflammation from CHIP to MDS. These observations suggest that while MDS loses overall HSPC-support and gains iMSCs, the inflammatory and HSPC-support programs may to some degree coexist within individual cells.

To further investigate the HSPC support function by MSCs, we predicted potential cell-cell interactions between individual MSC populations and the combined HSPC population using the NICHES tool ^89^. This analysis revealed significant changes in these interactions between control and MDS. In the controls, Adipo-CAR1 cells were identified as the major HSPC-interacting cell population (**Fig. 4G**), consistent with their known role in HSPC maintenance and the high HSPC support score (**Fig. 4A**). In CHIP, Adipo-CAR2 cells were the major interacting population, while in MDS the iMSC population became the predominant stromal cell type interacting with HSPCs (**Fig. 4G**). Notably, the iMSCs in MDS interacted significantly more with HSPCs than their iMSC counterparts in CHIP (OR ∼3.3; p-value >2.2e-16), supporting the idea that the iMSCs in CHIP and MDS we recovered from BM aspirates represent functionally distinct populations. This suggests that, while both conditions share an inflammatory signature, the iMSCs in MDS may be adapted to interact with and support HSPCs.

Together, these results suggest that the iMSC population in CHIP may differ from those in MDS or AML, both in terms of their HSPC support potential and response to inflammation.

### MDS blasts contribute to iMSC remodeling

Next, we sought to understand how MDS cells may contribute to remodeling of the stroma and emergence of iMSCs. First, we investigated the changes in HSPC populations across control, CHIP, and MDS donors. The most notable change was the significant reduction of Pre-B/Pro-B cells in both MDS and CHIP compared to controls (**Fig. 5A**), a finding consistent with reduced (mature) B-cells signatures seen in our NanoString data and the reduction of CD10^+^/CD19^+^ B cells with flow cytometry (**Fig. 1D, Fig. S1C**). However, no HSPC population was strongly increased in MDS or CHIP compared to the controls, with the exception of the CD14-low monocyte population that was similarly found to be increased in other blood malignancies such B-cell leukemia ^69^.

**Figure 5:**
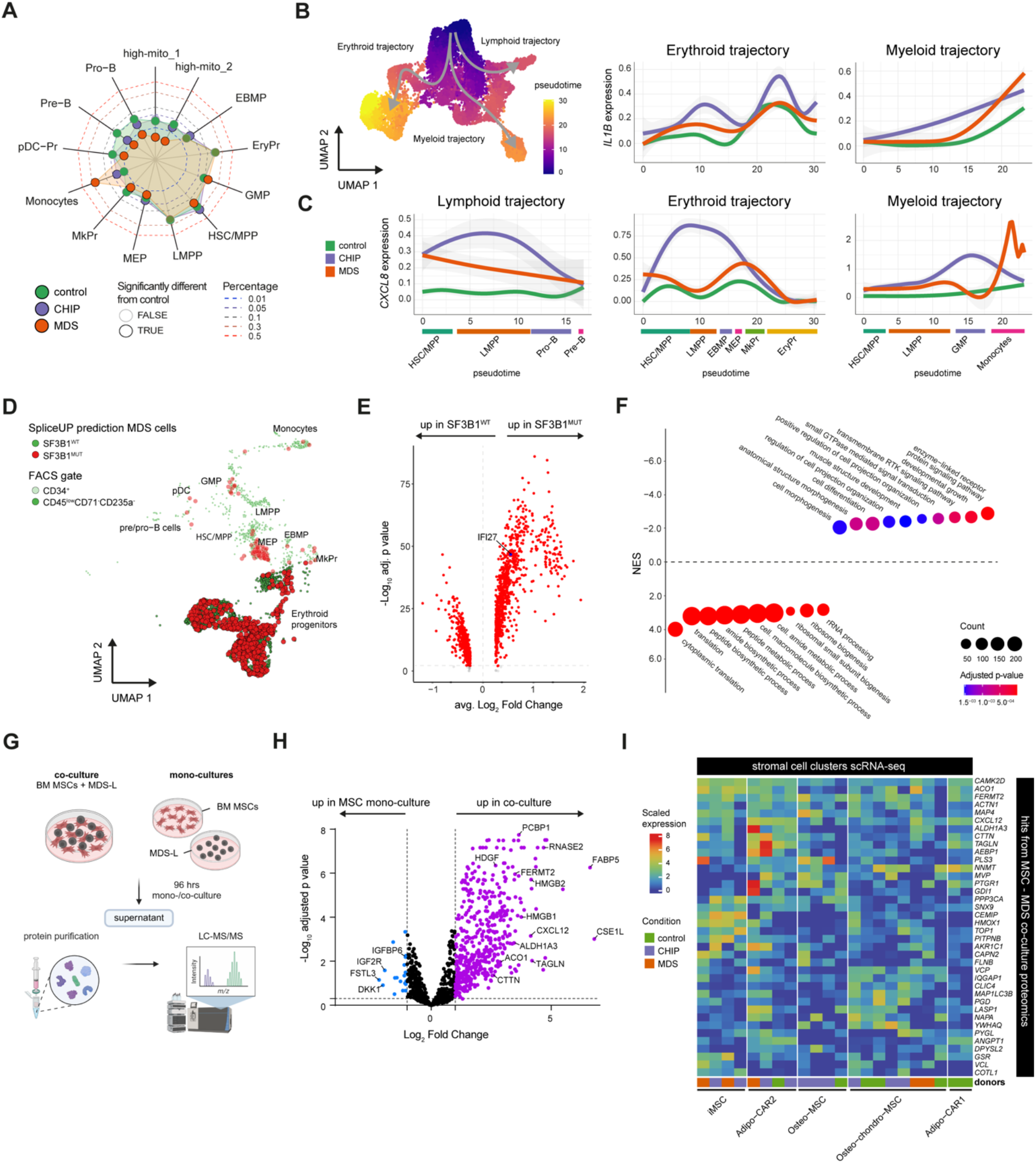
Characterization of HSPC and their effect on BM inflammation. **(A)** Spider plot displaying the relative abundance of cell types found in the HSPCs between different conditions control (green), CHIP (purple), and MDS (orange) donors. Cell type proportions are plotted to the radius after probit transformation, with a black outline on points indicating statistically significant differences compared to controls (see **Methods**). **(B)** UMAP of HSPC cells showing the pseudotime and schematic outline of main differentiation trajectories. **(C)** Smoothed expression of pro-inflammatory cytokine genes *CXCL8* and *IL1B* along the pseudotime in 3 major differentiation trajectories of HSPCs. **(D)** UMAP of HSPC cells showing the SpliceUp predictions of mutational status for SF3B1 gene. Mutation status and erythroid progenitor cells are highlighted in different colors. **(E)** Differential expression analysis results between SpliceUP-identified SF3B1^WT^ and SF3B1^MUT^ cells. 10% prediction-conditioned fallout (PCF) was used for the SpliceUP algorithm. Genes upregulated in the NanoString analysis between MDS and control samples (**Fig. 1**) are annotated with blue dots. Negative binomial test on single cells, with genes with adjusted p-value of < 0.01 annotated with red dots. **(F)** Gene Set Enrichment Analysis (GSEA) results with the Biological Process ontology from Gene Ontology database showing the top 20 significant sets enriched between SF3B1^WT^ and SF3B1^MUT^ erythroid progenitor cells predicted by SpliceUP. **(G)** Schematic summarizing the MDS-L and BM MSC co-culture experiment. MDS-L (5q) and BM MSC (hTERT) were cultured either separately or in a co-culture (ratio 1:1). After 96 hours supernatant was collected and subjected to protein purification and subsequent quantification using liquid chromatography-tandem mass spectrometry. **(H)** Differential protein abundance between BM MSC monoculture and BM MSC - MDS-L coculture. Labels show some of the significantly higher- or lower abundant proteins of interest. **(I)** Scaled average expression of genes matching proteins significantly upregulated in BM MSC - MDS-L co-culture compared to BM MSC mono-culture and specific to stromal cells compared to HSPCs in scRNA-seq data (adj. p-value <= 0.05, Wilcoxon rank-sum test) across stromal cell types. Expression values aggregated for each donor’s cells in each cell type

Aggregating expression across all HSPCs and quantifying differential expression between conditions, revealed a significant enrichment in pro-inflammatory pathways, such as TNFα via NF-κB, in MDS compared to control (**Fig. S7A**,**B**). Among the individual HSPC populations, this upregulation was observed in GMPs and LMPPs, suggesting these populations may drive the overall enrichment (**Fig. S7A**). A similar trend was also observed for all HSPCs in CHIP, but it did not reach statistical significance (**Fig. S7A**). To further investigate how differently committed HSPCs may contribute to inflammation, we used *Monocle3* to split the HSPCs into trajectories (erythroid, myeloid and lymphoid) ^90^ and quantified the expression of specific pro-inflammatory genes across pseudotime (**Fig. 5B**). We found that CHIP-derived HSPCs expressed high levels of pro-inflammatory genes CXC*L8* and *IL1β* across all lineages compared to controls. In MDS, HSPCs showed moderate increases in *CXCL8* expression across all lineages, with a pronounced increase in the monocyte population (**Fig. 5C**).

To determine whether the mutated HSPCs in MDS may drive inflammatory signaling, we applied our *SpliceUp* algorithm (*Kholmatov et al*., *in preparation*, **Methods**) to infer the SF3B1-mutational status of cells in MDS samples (n=3, **Fig. 5D**). Briefly, SpliceUp predicts mutated cells based on the presence of mis-splicing events characteristics for SF3B1-mutated cells. By aggregating data across multiple splice sites, SpliceUp circumvents the sparsity in single-cell transcriptomics data, reducing the number of cells without known mutation-status. This approach is more sensitive than directly looking for the mutant allele in RNA reads, though some SF3B1-mutant cells may still be misclassified as wild-type cells due to limited sensitivity.

We performed differential gene expression analysis between predicted SF3B1 wild-type (SF3B1^WT^) and mutant (SF3B1^MUT^) erythroid (progenitor) cells across 3 SF3B1-mutated MDS donors, including the mainly MDS-derived erythroid progenitor cells we “gained” during the stromal enrichment by FACS for the scRNA-seq (**Fig. S3I, Fig. 5D, Fig. S7C**). Overall, we identified 1132 genes differentially expressed between SF3B1^MUT^ and SF3B1^WT^ cells (**Fig. 5E**). The IFN-response gene *IFI27* was the only significantly upregulated gene in both scRNA-seq SF3B1^MUT^ cells and the MDS BM NanoString data (**Fig. 5D, Fig. S7D, Fig. 1F**), suggesting that the SF3B1^MUT^ HSPCs may have a limited direct contribution in driving the inflammatory milieu within the MDS. Moreover, GSEA analysis revealed that mRNA translation, peptide biosynthesis, and other metabolic processes were upregulated in SF3B1^MUT^ cells, while pathways related to cell differentiation and migration were enriched in SF3B1^WT^ cells (**Fig. 5F**). This suggests that SF3B1^MUT^ cells exhibit an adaptation towards increased cell proliferation in lieu of differentiation capacity, a hallmark of altered HSPC kinetics, which aligns with previous studies identifying this shift as a defining characteristic of MDS ^91,92^.

To further understand the direct effect of MDS blasts on MSCs, we established a co-culture system utilizing immortalized healthy BM MSCs (hTERT) and the MDS-L cell line, which carries mutations characteristic of 5q-MDS (*NRAS, CEBPA*, and *TP53*) ^93^, and has been previously linked to inflammation ^94^. MSC mono-cultures were compared to MSC-MDS-L co-cultures (1:1 ratio), and their secretome was analyzed by mass spectrometry (**Fig. 5G**). Following normalization and protein identification (**Fig. S8**), we identified 448 proteins that were significantly higher abundant in MSC-MDS co-culture vs MSC mono-culture (**Fig. 5H, Supplementary Table 9, Methods**). To pinpoint proteins secreted by MSCs (and not MDS cells) we used our single cell RNA-seq data and required that their RNA expression level was higher in stromal populations than HSPCs. Briefly, we averaged expression within each cell type and donor and selected the genes that were significantly higher expressed in stroma populations than HSPCs (adjusted p-value < 0.05, Wilcoxon Rank Sum Test; **Methods**), which resulted in 38 genes expressed in the stromal populations (**Fig. 5I**). Based on the expression profiles in our scRNA-seq data, the genes upregulated in MSCs upon co-culture are mostly present in our Adipo-CAR2 and MDS-iMSC populations. Among the proteins upregulated upon co-culture with MDS cells is the HSPC-support protein CXCL12, in line with the observation that iMSCs in MDS seem to have HSPC support capacity (**Fig. 4F**).

Overall, these results imply that MDS blasts contribute to building their own supportive niche, yet they seem not to directly contribute to the inflammatory milieu.

### MDS-specific IFN^+^ T cells interact with iMSC in MDS

To further understand what contributes to the inflammatory signature in CHIP and MDS BM, we investigated the T cell landscape in our data. Three distinct T cell populations were significantly less abundant in healthy controls: Heat Shock Protein (HSP)+ T cells were significantly enriched in CHIP and MDS, whereas IFN-responsive and cytotoxic CD8+ T_EMRA_/T_EFF_ cells were significantly enriched in MDS (**Fig. 6A**,**B**). HSP+ T cells have been previously described in the context of colorectal cancer, but a clear biological function has not yet been reported ^95^. Both IFN-responsive and cytotoxic CD8+ cells were recently associated with exacerbated inflammation in autoimmune diseases such rheumatoid arthritis ^96–98^

**Figure 6:**
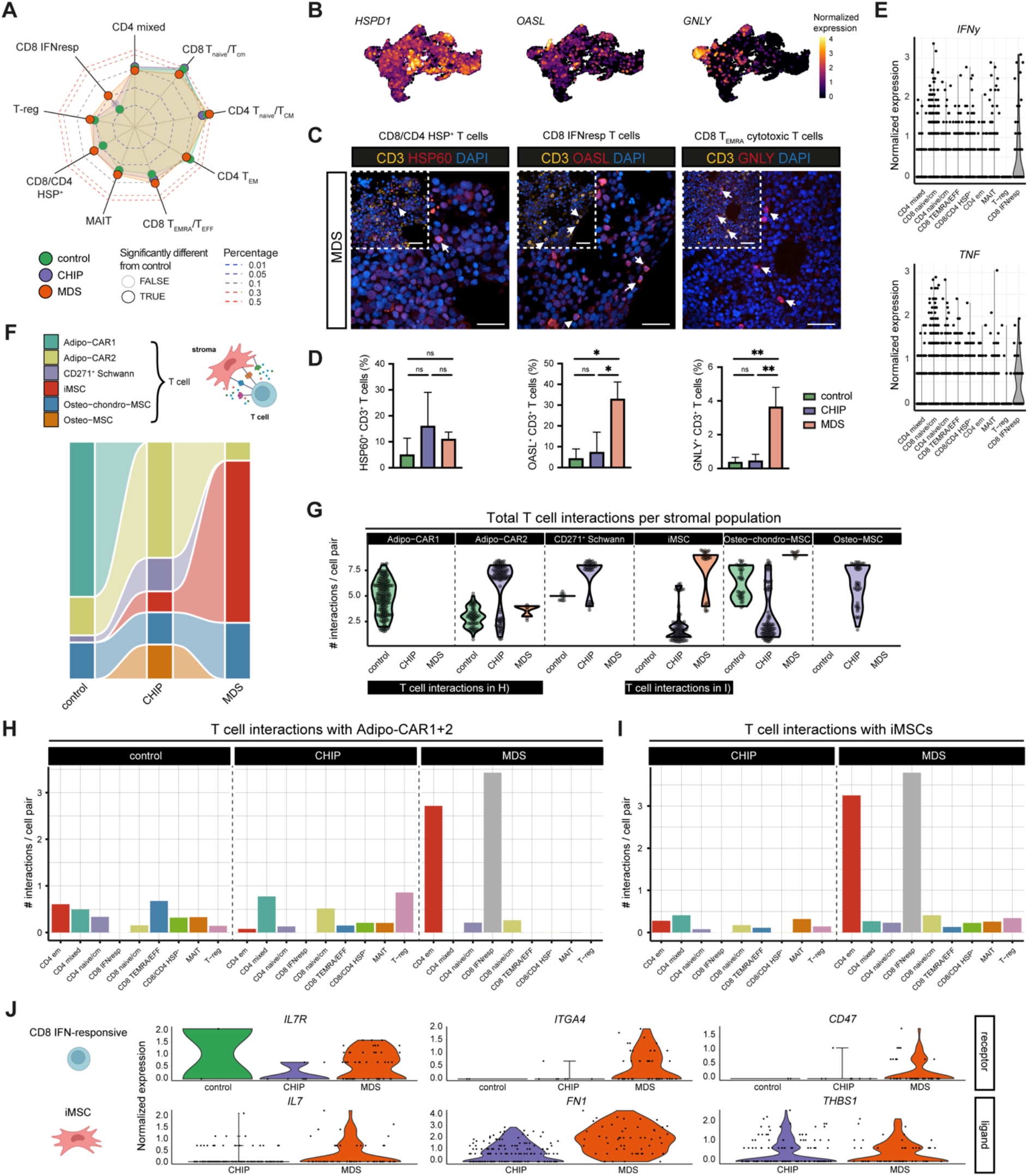
Characterization of the T cells in BM of healthy, CHIP and MDS. **(A)** Spider plot displaying the relative abundance of cell types found in the T cell population between different conditions control (green), CHIP (purple), and MDS (orange) donors. Cell type proportions are plotted to the radius after probit transformation, with a black outline on points indicating statistically significant differences compared to controls (see **Methods**). **(B)** UMAP plots showing the normalized expression of several genes highly expressed in populations that are enriched in CHIP and/or MDS conditions: CD8/CD4 HSP^+^, T_EMRA_ and CD8^+^ IFN-responsive. **(C)** Representative immunofluorescence images showing HSP60, OASL, and GNLY (red) representing the CD8/CD4 HSP^+^, CD8 IFN response T cells and CD8 T_EMRA_ respectively and CD3 (yellow) expression in MDS FFPE BM tissue sections. Insets highlight key regions of interest with distinct co-localization patterns. DAPI (blue) stains the nuclei. Scale bars are 25 μm. **(D)** Quantification of percent of cells expressing representative markers from B.) among all CD3+ cells in immunofluorescence imaging. Statistical significance: ns = not significant, * p < 0.05, one-way ANOVA (Tukey’s test for multiple comparison correction). **(E)** Violin plots of *IFNG* and *TNF* gene expression across T-cell subsets. Each dot represents the expression level of an individual cell. **(F)** Sankey diagram summarizing the proportions of inferred cell-cell interactions inferred by NICHES between T cells and different stromal cell types across control, CHIP, and MDS samples. **(G)** Average number of interactions between T cells and individual stromal cells per cell pair inferred by NICHES. **(H-I)** Total number of interactions between Adipo-CAR (H) or iMSC (I) stromal cells and individual T cell subtypes in different conditions. Numbers of interactions are normalized by the number of all possible interactions given the sizes of respective populations. **(J)** Expression of several ligand-receptor pairs found to be markers of interaction between iMSC and CD8^+^ IFN-responsive cells.

We validated the expansion of these inflammation-related T cell subsets in BM using immunofluorescence of CD3^+^ T cell compartment with previously reported markers also expressed in our scRNA data on a subset of our cohort. We used OASL for IFN-responsive T cells, GNLY for cytotoxic T T_EMRA_/T_EFF_ cells, HSP60 for HSP^+^ T cells ^99^; ^100^; ^101^ (**Fig. 6C, Fig. S9**). The OASL-expressing, IFN-responding and GNLY-expressing cytotoxic T_EMRA_/T_EFF_ cells showed a significant increase in MDS (**Fig. 6D**), confirming our scRNA-seq observations. In CHIP, the expansion of HSP^+^ T cells was visible but not statistically significant (**Fig. 6D**). These data confirmed that the T cells analyzed in the scRNA-seq data were not solely circulating T cells present in the BM vasculature.

IFN-responsive T cells showed elevated transcript levels of pro-inflammatory cytokines TNFα and IFNγ based on the normalized scRNA-seq data (**Fig. 6E)**. This suggests that IFN-responsive T cells may act self-or cross-activating in immune activation and thereby act as potent inflammation-driving populations.

Additionally, we have previously reported that a similar IFN-responsive T cell population in the BM of AML patients in remission after allogeneic stem cell transplantation were associated with imminent relapse ^65^, suggesting these T cells are not able to mount a response against AML cells. Comparing the expression signature upregulated in bulk BM in MDS vs control in our NanoString data revealed that T cells, and specifically IFN-responsive T cells, highly expressed genes involved in the inflammatory response (*IFIT1/2/3, TNF*; **Fig. 2I**). This implies these aberrant IFN-responsive CD8 T cells are key contributors to the inflammatory milieu in MDS.

By using the computational tool NICHES ^89^, we next investigated interactions between stromal populations and T cells, with stromal cells as senders and T cells as receivers. In both controls and CHIP samples, the Adipo-CAR1 and Adipo-CAR2 populations exhibited the most interactions with T cells respectively, whereas in MDS, iMSCs were the major interacting stromal population (**Fig. 6F**). Notably, MDS-iMSCs showed the highest number of interactions with T cells per MSC, suggesting extensive cross-talk between the iMSCs and T cells in MDS (**Fig. 6G**). In controls, Adipo-CARs interacted with all CD4/CD8 populations, in CHIP mostly with T-regs, CD4_naive/CM_, and CD4 mixed cells, while in MDS, their interactions were primarily with IFN-responsive T cells and CD4_EM_ cells (**Fig. 6H**). Notably, iMSCs in MDS had a very similar interaction profile to Adipo-CARs and strongly interacted with IFN-responsive T cells and CD4_EM_ cells (**Fig. 6I)**. This contrasts with iMSCs in CHIP, which displayed limited interactions with T cells, corroborating the observation that the iMSCs in CHIP and MDS represent two distinct functional populations (**Fig. 6I**).

We investigated the specific ligand-receptor interactions mediating the crosstalk between iMSCs and T cells in MDS. In particular, we identified several pairs enriched in MDS, including *IL7*, expressed by iMSCs, interacting with *IL7R* (*CD127*) on IFN-responsive T cells. Both the ligand (expressed by iMSCs) and receptor (on T cells) are upregulated in MDS compared to CHIP (**Fig 6J**). IL-7R expression can be induced by IFN signaling and IL-7 is a regulator of T cell differentiation and homeostasis ^102^. Other prominent ligand-receptor interaction included Fibronectin 1 (FN), secreted by iMSCs, interacting with Integrin 4A (ITG4A) on T cells, as well as Thrombospondin 1 (THBS1), an extracellular matrix glycoprotein produced by iMSC, interacting with CD47 on T cells (**Fig. 6J**). Notably, THBS1 was also among the proteins secreted by MSCs upon interaction with MDS blasts in our co-culture system (**Fig. 5H**), suggesting that the interaction of MSCs with MDS blasts may contribute to the recruitment of aberrant T cells to the BM, and specifically the iMSCs.

These results indicate that IFN-responsive T cells may contribute to the inflammatory signaling via TNFα and IFN-gamma secretion, while the blast-induced ECM remodeling of iMSCs in MDS may specifically attract inflammatory T cell populations.

### Aberrant cross-talk of MSCs and mutated HSPCs promotes vascular niche remodeling in MDS

Our integration of the targeted bulk RNA-seq data with healthy BM indicated elevated expression of inflammation-associated genes in endothelial cells in MDS (**Fig. 1G**). Previous studies have described expansion and remodeling of BM vasculature in myeloid malignancies, including MDS ^103–106^ increased expression of angiogenic factors including *VEGFA* in bulk BM^103–106^. Due to the limited availability of endothelial cells in human BM aspirates and the challenges in capturing them through FACS and scRNA-seq, we assessed the angiogenic potential of BM cell types: HSPCs, T cells, and stromal populations (**Fig. 7A, Fig. S10A-C**). We found that the Adipo-CAR and iMSC populations were primarily responsible for angiogenesis-related condition-dependent changes (**Fig. 7A, Fig. S10C)**, while other cell populations showed minimal angiogenic potential (**Fig. S10A**,**B**). This is in line with the reported localization of Adipo-CARs in vascular BM niches in mouse and human BM ^47,107–109^. Specifically, Adipo-CARs in MDS patients upregulated several known secreted vasculature-remodeling factors, including, *VEGFA* and *FSTL1*, which were also secreted by iMSCs both in CHIP and MDS (**Fig. 7B**). These results suggest that on top of increased inflammation and reduced HSPC-support, the remodeled stroma in MDS may reshape the BM vasculature. We leveraged immunofluorescence imaging on FFPE BM biopsies to investigate this increased angiogenic potential within the MDS niche on the arteriolar and sinusoidal blood vessels (**Fig. 7C**). We quantified the microvasculature density (MVD) for both arteriolar (UEA1^+^CD105^-^) and sinusoidal (UEA1^-^CD105^+^) vasculature, which revealed a significant increase in MVD for both structures in MDS compared to control and CHIP (**Fig. 7D**,**E**). Additionally, we observed a reduction in the average distance between sinusoidal and arteriolar vessels in MDS compared to CHIP and control (**Fig. 7F**).

**Figure 7:**
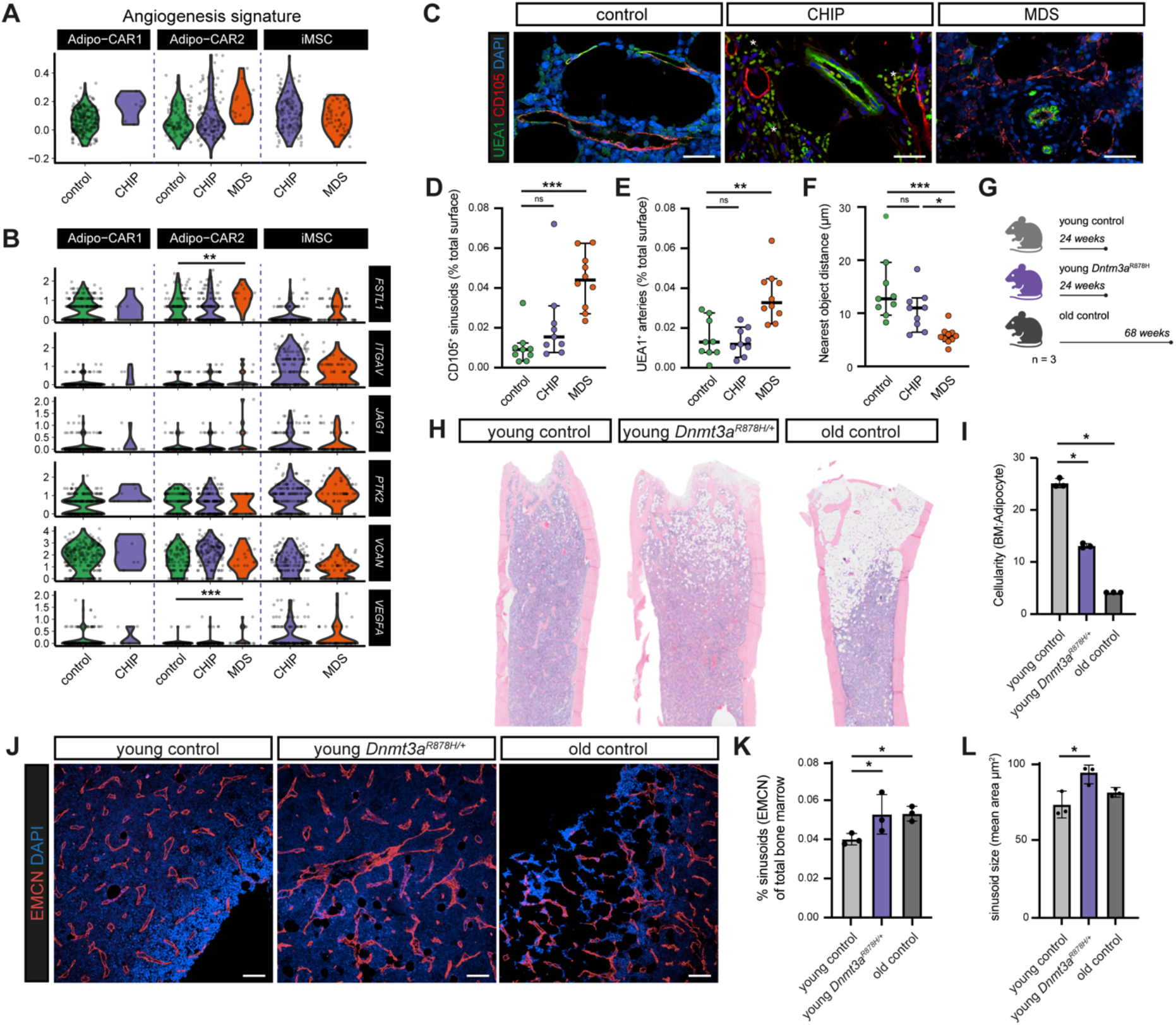
Assessment of vascular alterations in human CHIP, MDS, and CHIP DNMT3A mouse model. **(A)** Violin plots showing the distribution of angiogenesis signature scores across the stromal cell types comparing control, CHIP and MDS. **(B)** Violin plots showing the expression of most representative genes of angiogenesis signature across the stromal cell types comparing control, CHIP and MDS. Statistical significance: ** p < 0.01, *** p < 0.005, Wilcoxon rank-sum test. **(C)** Immunofluorescence staining of UEA1 (all vasculature) and CD105 (sinusoids) on FFPE BM tissue sections. Representative images showing UEA1 (Green), CD105 (Red), and DAPI (blue) staining in control, CHIP, and MDS FFPE BM sections. Asterisks are highlighting autofluorescence from erythrocytes. Scale bars are 25 μm. **(D-E)** Quantification of sinusoids (D) and arterioles/arteries (E), showing the relative abundance of CD105^+^ sinusoids (D) and CD105^-^UEA1^+^ arteries (E) in control (green), CHIP (purple), and MDS (orange) samples. MDS samples exhibit a significant increase of both vasculature types compared to controls. Statistical significance: ns = non-significant, * p < 0.05, ** p < 0.005, *** p < 0.001, Kruskal-Wallis test. **(F)** Scatter plot depicting the nearest distance of the vasculature (in um) in control, CHIP, and MDS samples. Shortened distances in MDS indicate disrupted vasculature organization. Statistical significance: * q < 0.0001, one-way ANOVA (FDR for multiple comparison correction). **(G)** Schematic summarizing the three groups in the CHIP *Dnmt3a*-mutant mouse study: femurs were collected from young control (24 weeks), young *Dnmt3a*^R878H^ (24 weeks), and old control (68 weeks). 3 mice per group were analyzed (n=3). **(H)** Giemsa staining from young control, young *Dnmt3a*^R878H^, and old control mouse femurs, illustrating reduced cellularity in young *Dnmt3a*^R878H^ and old mice. **(I)** Bar plot showing the cellularity (BM/adipocyte ratio) in the conditions. Both young *Dnmt3a*^R878H^ and old control mice display significantly reduced cellularity compared to young controls. **(J)** Representative immunofluorescence images of EMCN (red) and DAPI (blue) staining in FFPE femur sections from young control, young *Dnmt3a*^R878H^, and old control mice. Scale bars are 100 μm. **(K)** Quantification of the total sinusoid area in the femur bone. Statistical significance: * adjusted P-value < 0.05, two-way ANOVA (Tukey’s test for multiple comparison correction). **(L)** Quantification of the mean area of the sinusoidal vasculature (sinusoid size). *Dnmt3a*^R878H^ mice show in addition to increased sinusoidal also a widening of the sinusoidal structures. Statistical significance: * q < 0.05, one-way ANOVA (FDR for multiple comparison correction).

Despite the presence of iMSCs with angiogenic potential within CHIP donors (**Fig. 7A**,**B**), no significant vascular changes were observed (**Fig. 7C-F**). Given the correlation between osteoporosis risk and a higher VAF% of the CHIP mutation *DNMT3A*^R882H^ (>10%) ^110,111^, we further explored the yet uncharted impact of VAF on vasculature remodeling. We utilized a CHIP mouse model carrying the *Dnmt3a*^R878H^ mutation, which is homologous to a common human *DNMT3A*^R882H^ CHIP mutation, where the majority of hematopoietic cells carry this mutation ^112^. We analyzed the general BM morphology and vasculature within femurs of young control mice (24 weeks), young *Dnmt3a*^R878H/+^ mice (24 weeks), and old control mice (68 weeks) (**Fig. 7G**). First, we showed that *Dnmt3a*^R878H/+^ decreased the overall cellularity in BM, driven by the expansion of adipocytes particularly in the femoral head (**Fig. 7H**,**I, Supplementary Table 10**). Next, we observed that young *Dnmt3a*-mutant mice showed increased sinusoidal vessel density (**Fig. 7J**,**L**) and general morphological alterations including increased dilation of the sinusoidal vasculature (**Fig. 7K**), which is in line with our observations in human MDS (**Fig. 7C-F**). Arterioles and arteries did not show any significant differences between conditions (**Fig. S10D**). This suggests the higher burden of mutant cells in CHIP may drive BM niche architecture remodeling, however, additional studies in human individuals with a higher CHIP VAF will be needed to validate this.

Altogether, these results suggest that mutant HSPCs can significantly alter the BM niche vasculature, potentially via stromal remodeling. These findings provide new insights into how CHIP may prime the BM microenvironment for malignant transformation.

## Discussion

Our study presents a comprehensive dataset characterizing the stromal and T cell landscape of human bone marrow (BM) in CHIP and MDS. Our findings emphasize the dynamic changes in the microenvironment in response to mutated hematopoietic cells and their role in fueling inflammation.

Previous studies based on global gene expression analysis from MSC-derived cultures and bulk sequencing hinted at an inflammatory-primed state in the context of aging, chronic bone-related diseases, and MDS ^113–115 42,116,117^. Our study highlights the presence of a distinct inflammatory MSC (iMSC) population that is expanded in CHIP and MDS but absent in aged control BM. Similar iMSCs have recently been reported in other single cell studies on BM malignancies, including AML ^85^ and Multiple Myeloma (MM) ^76^. Our comparative analysis of these previously identified inflammatory/responsive MSC populations demonstrated a high resemblance of their inflammatory gene expression profiles toward our characterized CHIP/MDS iMSCs. In our study, iMSCs were identified within a homogeneous cohort of low-risk and mostly spliceosome-factor mutated MDS, allowing for a comprehensive interpatient comparison with a similar mutational spectrum. The discovery of iMSCs was also confirmed in a parallel study by Chen et al. (co-submitted) in a more mutational heterogeneous MDS cohort. The absence or low abundance of this iMSC subset in healthy controls across all datasets underscores its potential role in disease progression. Additionally, inflammatory stimuli in MSC cell lines have been shown to improve bone regeneration when transplanted in mice ^118^. This may suggest that the observed expansion of osteo-lineage MSCs in our dataset, balanced with the reduction of the adipo-lineage, in CHIP and MDS is driven by the inflammatory milieu in these conditions. Further characterization and functional investigation may reveal a similar role of iMSCs in CHIP/MDS.

In our study, iMSCs display elevated CD44 expression in CHIP and MDS and are characterized by high ECM production. Pro-inflammatory cytokines and growth factors, such as TNFa, IL-6, and IL-1β, have been shown to induce *CD44* promoter activity through several transcription factors, including SP1, EGR1, and NF-κB ^119–124^. Moreover, involvement of CD44 in cell-cell interactions is linked in modulating inflammation and tumorigenesis ^125^. In parallel, we identified a specific upregulation of IL-1R1 within the iMSC population, known as a specific inflammatory marker of cancer-associated fibroblasts (CAFs) in solid tumors ^126,127^. A recent study identified a subset of IL-1R1-high CAFs in colorectal cancer with enhanced motility and inflammatory response that drive tumor development by suppressing T cell and macrophage functions ^128^. Taken together, the specific upregulation of CD44 and IL-1R1 within the iMSC population may point to a shared inflammatory gene program between stromal cells in hematological malignancies and CAFs in solid tumors ^83,129^.

While both CHIP and MDS exhibit an inflammatory stromal response, our comparative analysis suggests the iMSC population are not the same in CHIP and MDS. Specifically, in CHIP the inflammatory and HSPC-support phenotypes are represented by two separate MSC populations, whereas in MDS the iMSC subset shows a combined HSPC-support/inflammatory phenotype. A recent study by Jann and colleagues ^130^ examined the stromal support of hematopoiesis in an MDS-xenograft mouse model and cross-referenced their findings with a small, FACS-enriched MSC population from a mixed MDS patient cohort analyzed at single-cell level. In the PDX mouse model, they found that CAR cells (CXCL12-abundant reticular cells) exposed to MDS cells had increased expression of key hematopoietic support factors, including *Cxcl12, Kitl*, and *Il7*. In the human BM MSCs, an elevated IFNα and IFNy response was noted in MDS compared to healthy controls, along with higher CXCL12 expression in particularly the “common CAR” population. However, this increased hematopoietic support signature was not consistent across all donors. Our findings align with this, as iMSCs in LR-MDS expressed higher levels of hematopoietic support genes than in CHIP. Additionally, LR-MDS cells were able to upregulate CXCL12 in co-cultured MSCs, though the overall hematopoietic support in MDS remains lower than in controls. Interestingly, AML-associated stromal cells are described to have a reduced expression of hematopoietic support factors (e.g., CXCL12, LEPR, KITLG) ^85^, suggesting that AML (leukemic stem cells) may thrive in a support-deprived environment. Potential differences in stromal response between AML and MDS might be due to the dependency on the stemness status of the malignancy. AML, with its circulating immature blasts, can proliferate despite low HSPC support cues, while MDS blasts still rely on the BM niche. Potentially, leukemic stem cells and blasts from different myeloid malignancies interact with MSCs in different BM niches. Moreover, the increased expression of angiogenic factors by both Adipo-CAR and iMSC correlated with expansion of microvasculature in MDS, suggesting a greater contact surface between vascular walls and HSPCs. This increased contact surface may contribute to increased exposure of mutated HSPCs to different vascular and stromal cues. In addition, sinusoidal vessels in MDS showed a disorganized structure and signs of increased membrane permeability, which may contribute to impaired oxygen and nutrient delivery, further destabilizing the BM microenvironment. This may lead to hyperactivation of malignant HSPCs and increased proliferation, supported by recent reports linking high vascular permeability to an increased HSPC activation from quiescence ^131^ and the egress of HSPCs from the BM to peripheral circulation in AML ^132^. These changes may contribute to HSPC dysfunction and enhanced proliferation of mutant cells, driving disease progression. Future research should focus on understanding how hematopoietic and angiogenic support varies across specialized niches of the human BM, for example using advanced spatial mapping techniques.

In addition to the progressive dysfunction of the hematopoietic MSC niche, our data demonstrated that inflammation-induced changes in CHIP and MDS affect the T cell landscape in BM. Unlike more advanced hematological malignancies such as AML and MM ^133,134^, where significant T cell alterations are observed, only a few T cell states were impacted in CHIP and low-risk MDS. In CHIP, a CD4^+^/CD8^+^ subset expressing high levels of heat shock protein (HSP) family members was enriched and may represent a stress response of naive T cells to the inflammatory environment created by (mutated) HSPCs. Although not well characterized, these HSP^+^ T cells have recently been proposed to act as immunoregulators of chronic inflammation ^135^ and linked to malignant progression in glioma ^136^. Further investigation and experimental validation are warranted to clarify their role in CHIP emergence and progression.

In MDS, accumulation of IFN-response T cell subsets might represent a locally activated immune state, possibly driven by TCR-triggered IFNγ or induced by interferons directly; as shown in previous large pan-cancer atlases of T cell activation signatures ^63,137^. The presence of IFN-responsive T cells in MDS may arise as a reaction to the inflammatory signals from HSPCs such IL-8, similar to those seen in pathogen responses ^138,139^. Our inferred cell-cell crosstalk predicted an almost exclusive interaction of IFN-responsive T cells with iMSCs via the IL-7/IL-7R and ITGA4/FN1 axes, which may fuel further inflammation back to the BM niche. IL-7, a regulator of T cell differentiation and homeostasis ^102^, could drive the expansion and survival of T cells, with IFN signaling possibly upregulating IL-7R expression in IFN-responsive T cells, making them more susceptible to IL-7 secreted by iMSCs. Additionally, Fibronectin 1 (FN1) secreted by iMSC could recruit T cells (expressing ITGA4) and other immune cells to the BM. Lastly, the expanded CD8^+^ T_EMRA_ cells might also harbor tumor-specific TCRs, which is consistent with the notion of a systemic immune response ^140,141^. Thus, there is a need for increasing research efforts to deepen our understanding of T cell biology and implication in CHIP and MDS, which could bring novel approaches to treatment.

Both CHIP and MDS exhibited a primed inflammatory status that impacted the T cell landscape and possibly other immune cells. This is supported by the upregulation of pro-inflammatory cytokines (IL-1β, CXCL8) in HSPCs and chemokines in iMSCs (CCL2, MCP-1), which can recruit monocytes ^142,143^ and may promote their differentiation into pro-inflammatory macrophages, as seen in chronic bone diseases ^79,144,145^. This suggests that the stromal niche can exacerbate immune dysregulation in MDS, but further investigation into monocytes and activated macrophages is needed. Accordingly, we observed unexpectedly non-classical monocytes expressing tumor-associated inflammatory factors (ADORA2A, S100A8), resembling myeloid-derived suppressor cells (MDSCs) ^146,147^ which have been implicated with immune dysregulation in MDS ^148^. ARG1, an MDSC marker ^149^, was upregulated, suggesting a chronic exposure to inflammatory cytokines and myeloid growth factors (IL-6, IL-1β, S100A8/9) ^150^, potentially driven by the expanded iMSC population. This could promote immune suppression and further fuel disease progression.

Considering the unusual interplay between mutated HSPCs and emerging inflammatory populations in the stroma and T cell compartments, it is plausible that the size of the clone or the variant allele frequency (VAF) of CHIP mutations in the HSPC compartment influences niche remodeling and the subsequent development of MDS. Previous studies have categorized CHIP carriers into low- and high-VAF groups when the clone size exceeds 10%, linking high-VAF CHIP to an increased risk of osteoporosis, cardiovascular diseases, and various solid and hematological cancers ^33,110,151,152^. It is not surprising that systemic inflammatory diseases are associated with VAF, given that mutated HSPCs have been shown to promote pro-inflammatory signaling. The stroma remodeling effects observed in this study may also depend on clone size, as they are detectable in low-VAF CHIP via scRNA-seq but not in tissue staining. In comparison, apparent BM architecture remodeling and vascular reshape is detected in the MDS cohort, where the VAF is approximately 40%. Similar features of vascular defects become noticeable in a CHIP mouse model in which most hematopoietic cells carry a *Dnmt3a* mutation. While MDS pathology involves factors beyond the expansion of mutant HSPC clones and cannot be equated with true high-VAF CHIP samples, there are evident common stromal alterations in both conditions that highlight the bone marrow microenvironment as a potential mediator in the emergence of MDS. In conclusion, our findings highlight a shared inflammatory response across myeloid malignancies, suggesting that chronic inflammatory stimuli shape the BM microenvironment in both CHIP and MDS, albeit with distinct alterations in their stromal architecture and T cell landscapes. This research underscores the need for further exploration of the mechanisms underlying these inflammatory subsets and immune responses, which may offer new therapeutic targets and strategies for managing these conditions in future studies.

## Materials and Methods

### Patient cohort and tissue material processing

Overall, BM samples from 32 MDS patients, 17 CHIP donors, and 35 healthy (control) donors were analyzed within this study. Human BM aspirate and hip and femur trephine core biopsies were obtained from consenting donors enrolled in the BoHemE study (NCT02867085). MDS, CHIP, and control samples had approval from local ethics committees at the University Hospitals Dresden and Leipzig under broad research informed consent with unspecified future to be used as part of the MDS registry (EK289112008) or BoHemE study (EK393092016, 137/19-lk). A detailed overview of the patient cohort can be found in **Supplementary Table 1**. Aspirates were processed by density gradient centrifugation to deplete erythrocytes and were stored in freezing medium (90% FBS + 10% DMSO) in liquid nitrogen. Trephine cores were fixed in 4% PFA for 24 hrs, then transferred to PBS with sodium azide (0.3%) until further processing. Samples were decalcified for 48 hrs using Osteosoft (Merck KGaA) at 37°C and embedded in paraffin (FFPE).

### Animal models

Femurs from control and DNMT3A-R878H mutant mice were obtained from The Jackson Laboratory, USA. Briefly, *Dnmt3a*^*fl-R878H/+*^ mice (JAX:032289) were crossed with B6.Cg-Tg(Mx1-cre)1Cgn/J mice (JAX:003556, referred to as Mx-Cre). In all experiments, control (+/+) mice carried a single copy of the Mx-Cre allele. To induce Mx-Cre, mice were injected intraperitoneally with 15 mg/kg high molecular weight polyinosinic-polycytidylic acid (polyI:C) (InvivoGen) once every other day for a total of five injections. In all experiments, mice were used >4 weeks after polyI:C administration. Following the Jackson Laboratory recommendations for aging stages, mice up to 30 weeks (6-7 months) were considered “young”, while mice 68 weeks (15-16 months) and older were considered “old”. All experimental mice were sacrificed between 26 and 86 weeks of age for phenotyping and tissue collection. Femurs were fixed in 4% PFA at 4°C overnight and stored in PBS + 0.3% sodium azide. After decalcification, bones were paraffin embedded. The Jackson Laboratory Institutional Animal Care and Use Committee approved all experiments.

### Flow cytometry of BM aspirates

#### Diagnostic flow cytometry

Diagnostic flow cytometry was performed within 24 hrs after BM aspiration. Staining, acquisition, and analysis were performed as described previously ^153^. Prior to antibody staining, erythrocytes were removed by lysis for 10 min at room temperature using BD Pharm-Lyse (1:10 dilution with distilled water; BD Biosciences), followed by two washing steps with PBS. For surface labeling, the cells were incubated with one of the five 8-color antibody panels (see **Supplementary Table 3** for antibody panels) for 15 min at room temperature in the dark. The five panels allowed a comprehensive analysis of the BM aspirates as proposed in the guidelines of the iMDSFlow ^154^. Subsequently, cells were washed twice and resuspended in PBS. Samples were stored at 4°C and measured within 1 hr. Samples were acquired on a FACS Canto II (BD Biosciences). At least 200,000 events were acquired per sample. For data analysis, a hierarchical gating strategy according to the iMDSFlow ^154^ was applied: (1) exclusion of doublets (FSC-A vs. FSC-H) and (2) of debris (FSC vs. SSC), (3) gating of CD45^dim/+^ leukocytes (SSC vs. CD45).

#### Inflammatory MSC flow cytometry

BM aspirates from the curated BoHemE study (NCT02867085) were thawed in 100% FCS + 100 μg/ml DNAse, centrifuged at 300g for 5 min and suspended in DMEM + 10% FCS + 100 μg/ml DNAse. When necessary, cell pellets were treated with 1X Red Blood Cell Lysis (Invitrogen) for 10 min at room temperature. Cells were washed with 2% FCS/PBS. Live/dead staining (Zombie Aqua, Biolegend) was performed in PBS, followed by antibody staining for 30 min at 4°C (see **Supplementary Table 3** for antibody panel and dilutions). Stained cells were analyzed with a FACSAria™ Fusion (BD Biosciences).

### Immunohistochemistry and histopathological stainings of FFPE bone marrow sections

Serial sections of human and mouse FFPE BM biopsies were prepared at 3-5 μm thickness on coated microscope slides (Dako FLEX, Agilent) and processed for immunohistochemistry/immunofluorescent-based (IHC-IF). Deparaffinization (30 min), rehydration (2x 10 min), and antigen retrieval (10 min) were performed by heating sections in Trilogy™ buffer (Cell Marque, Millipore Sigma) at 105°C and 1.2 bar for 10 min in a pressure cooker. Sections were permeabilized with 0.25% Triton-X in PBS for 10 min, then blocked for 30 min with 5% normal donkey serum (Jackson ImmunoResearch) in PBS-Tween20 (0.05%). Primary antibodies were incubated at room temperature for 1 hr in 1% normal donkey serum PBS-Triton-X100 (0.2%). Samples were washed twice with 1% donkey serum PBS-Tween20 (0.05%) for 5 min, then incubated with secondary antibody for 1 hr at RT. Samples were washed again with 1% donkey serum PBS-Tween20 (0.05%) for 5 min, then incubated for 3 min with the TrueView (Vector Laboratories) quenching reagent to reduce tissue autofluorescence. Slides were washed with PBS for 5 min, mounted with DAPI-containing mounting medium (Abcam), and imaged. All used antibodies in IHC-IF are detailed in **Supplementary Table 3**. Additional FFPE sections of BM biopsies were processed for standard pathology diagnosis with Giemsa staining by the Core Facility Biobank of the UMC Mainz.

### Image data acquisition and analysis

IHC-IF stained sections were imaged for semi-quantitative analysis on an Opera Phenix (Perkin Elmer) high-content screening system with 40x objective (water, NA 1.1). Image analysis was done using Harmony 4.8 (Perkin Elmer), detailed workflows for image analysis can be found in **Fig. S4**.

Giemsa-stained sections were imaged using an EVOS M5000 microscope with 4x objective (air, NA 0.13). Captured images were analyzed for cellularity analysis using the Weka-Segmentation plugin for Fiji; for each condition, one representative image was used as training data ^155^. For each sample, three different fields of views were analyzed and the resulting probability maps were checked for successful segmentation. The detailed workflow can be found in **Fig. S4**. For the Giemsa staining of mouse femurs, cellularity analysis was conducted using QuPath software with the MarrowQuant plugin ^156^.

### Cell lines and co-culture conditions

The human cell line hTERT-MSC was kindly provided by Dr. Charles Mullighan (St. Jude Children’s Research Hospital) and cultured under sterile conditions in aMEM (Gibco) supplemented with 7% human Platelet Lysate (hPL, Macopharma) and Penicillin/Streptomycin (Gibco). The MDS-L cell line (derived from a RARS patient and SF3B1 WT ^157^) were kindly gifted by Dr. Kaoru Tohyama (Kawasaki Medical School) and cultured under sterile conditions in RPMI 1640 containing Glutamax (Gibco), supplemented with 10% FBS (Gibco), 40 ng/ml GM-CSF (Peprotech), and 1x Penicillin/Streptomycin. All cell lines were routinely tested for mycoplasma and authenticated by Single Nucleotide Polymorphism (SNP)-profiling (Multiplexion, Heidelberg, Germany). All cell lines were maintained up to 70-80% confluence in a HERAcell incubator (Thermo Fisher Scientific) at 37°C, 5% CO_2_ in air, and a humidity of ∼90%.

For the proteomics secretome analysis, *in vitro* co-cultures of hTERT-MSC and MDS-L were established by seeding 200,000 hTERT-MSCs overnight in a 6-well plate. After 24 hrs, culture medium was aspirated and 200,000 MDS-L cells, washed 5 times in PBS to remove residual FBS, were added in medium containing aMEM + 7% hPL. MSC-MDS co-cultures with equal numbers per cell type were defined as ratio 1:1, with additional conditions at ratio 10:1 and 1:10. Mono-cultures of both cell lines (ratio 1:0 and 0:1) were used as controls. After cultivation of 96 hrs, the conditioned supernatant was collected and centrifuged at 300g for 5 min, then the supernatant at 10,000g for 5 min to exclude any cell debris, and stored at -20°C until further use for proteomics analysis (see below).

### Mass spectrometry of MDS-MSC co-culture secretome

Supernatants of mono-cultured MSCs, mono-cultured MDS-L, and co-cultures of both cell types (in 10:1, 1:1, and 1:10 ratios) were harvested in quadruplicates and prepared for mass spectrometric analysis using label-free quantification (LFQ) with a slightly modified single-pot, solid-phase-enhanced sample-preparation (SP3) protocol, as previously described ^158^. Briefly, 350 μl of culture supernatant was mixed with 38.9 μl of a 1M aqueous solution of 2-[4-(2-hydroxyethyl)piperazin-1-yl]ethanesulfonic acid (pH 7.5), 8.75 μl bead stock (1:1 mix of Sera-Mag SpeedBeads GE45152105050250 and GE65152105050250 at 20 μg/μl, Merck) and 875 μl acetonitrile. Beads were washed, proteins reduced with dithiothreitol, alkylated by iodoacetamide, and finally digested by adding 2x 0.05 μg Trypsin in 50 mM ammonium hydrogen carbonate. Peptides were dried down in a vacuum concentrator and resuspended in 0.1% trifluoroacetic acid in water, and half of the samples were analyzed by liquid-chromatography coupled mass spectrometry. Here, peptides were separated for 1 hr on C18 material by an Ultimate 3000 Rapid Separation Liquid Chromatography system (RSLC, Thermo Fisher Scientific) as previously described ^159^, sprayed via a nano-electrospray source into an online coupled QExactive plus mass spectrometer (Thermo Fisher Scientific) operated in positive, data-dependent mode. First, survey scans were recorded (resolution: 140,000, target value advanced gain control: 3,000,000, maximum ion time: 50 ms, scan range 200-2000 m/z, profile mode). Second, up to 20 2- and 3-fold charged precursors were isolated by the quadrupole of the instrument (isolation window: 4 m/z), fragmented by higher-energy collisional dissociation (normalized collision energy: 30), and analyzed in the orbitrap (resolution: 17,500, target value advanced gain control: 10,000, maximum ion time: 50 ms, available scan range 200-2000 m/z, centroid mode). Already fragmented precursors were excluded from the analysis for the next 10 seconds.

Protein identification and quantification were enabled by *MaxQuant v*.*1*.*6*.*17*.*0* (Max Planck Institute of Biochemistry, Planegg, Germany) with standard parameters if not stated otherwise. Proteins sequences were provided by the UniProtKB (75777 homo sapiens entries, downloaded on 27th January 2021, UP000005640), the match-between runs function as well as label-free quantification enabled. Quantitative data was processed by *Perseus* (Max Planck Institute of Biochemistry, Planegg, Germany) and proteins were annotated with Gene Ontology (GO) and Kyoto Encyclopedia of Genes and Genomes (KEGG) enrichment terms provided by *Perseus* ^160^ as well as prediction of the mode of secretion by *OutCyte* ^161^. Prior to differential analysis, LFQ data was pre-filtered by removing potential contaminant sequences, reverse sequences and proteins that are identified by a single peptide. In addition, proteins with a high proportion of missing values are removed with less than three valid quantitative values (biological replicate) in at least one sample group (condition). Proteins with missing values were retained in the analysis based on a dynamic exclusion strategy ^162^: if it met the requirement of at least four valid values in at least one sample group comparing mono-versus 1:10 or 10:1 co-cultures respectively; or at least three valid values in one sample group when comparing both mono-cultures.

After pre-filtering, all LFQ intensities were transformed to a Log2 scale and replicates were grouped by conditions. Missing values were imputed using the ‘Gaussian random sample’ method with single-value replacements, which uses random draws from a Gaussian distribution left-shifted by 1.8 StDev (standard deviation) with a width of 0.3 ^162^.

Finally, protein-wise linear models combined with empirical Bayes statistics were used for the differential expression analyses (all conditions) by using R-based *LFQ-Analyst* with Bioconductor package *limma* ^163^, whereby the adjusted P value cutoff was set at 0.05. For significance analysis, the Benjamini-Hochberg method of False Discovery Rate (FDR) correction was used to account for multiple testing.

### NanoString nCounter gene expression analysis

Total mRNA was isolated using AllPrep DNA/RNA Mini Kit (Qiagen, Hilden, Germany) following the manufacturer’s instructions. 150-300 ng RNA was extracted per sample from 36 whole BM aspirates (12 MDS patients, 12 CHIP donors, and 12 controls). The expression of 1255 unique genes were analyzed in the NanoString nCounter Pro Analysis System using the PanCancer Immune profiling panel and Immune Exhaustion panel (NanoString Technologies, Amsterdam, Netherlands). The *nSolver* software package and *nSolver Advanced Analysis* module (NanoString Technologies) were used to evaluate the determined transcript counts. Quality-checked raw data was normalized utilizing the geometric mean of the housekeeping reference genes and the code sets’ internal positive controls were used to exclude samples that are outliers. Statistics were calculated in *R* based on normalized and log2-scaled counts applying an empirical Bayes moderated t-statistics tests using *limma (v*.*3*.*58*.*1)* ^164^ and corrected for multiple testing adopting the Benjamini-Hochberg procedure. Genes were determined differentially expressed with FDR < 0.05. For comparing CHIP and control samples, log2-fold changes were ranked based on their magnitude without applying an FDR cut-off and determined as differential with an absolute fold change > 0.5. Gene Ontology (GO) and Kyoto Encyclopedia of Genes and Genomes (KEGG) enrichment of differentially expressed genes was done with *clusterProfiler v*.*4*.*10*.*1* ^165^ using all genes measured in the same panel as background. P-values of GO and KEGG enrichment results were corrected for multiple testing applying Benjamini-Hochberg procedure and determined significant with an FDR < 0.05. Top 10 significant GO- and KEGG terms were selected by magnitude of FDR for plotting. Additionally, significantly changed genes were submitted to Gene Set Enrichment Analysis (GSEA) against the Hallmark and the C6 gene sets ^166,167^.

### Healthy BM single cell atlas integration with NanoString data

Publicly available healthy BM scRNA-seq dataset from ^47^ was acquired. Expression of genes found to be significantly up- or down-regulated between the MDS and control samples in NanoString analysis (both panels used) at FDR <= 0.05 was aggregated based on either the cell type annotation provided by the authors (**Fig. S2H**) or a simplified annotation grouping related cell types **(Fig. 1G**) using Seurat’s *AverageExpression* function and visualized using the ComplexHeatmap R package ^168^.

### Cell isolation/processing of hematopoietic and non-hematopoietic cells for scRNA-seq

BM aspirates from the curated BoHemE study (NCT02867085) were thawed in 100% FCS

+ 100 μg/ml DNAse, centrifuged at 300g for 5 min and suspended in DMEM + 10% FCS + 100 μg/ml DNAse. When necessary, cell pellets were treated with 1X Red Blood Cell Lysis (Invitrogen) for 10 min at room temperature. Cells were washed with 2% FCS/PBS. Live/dead staining (FV780, Thermo Fisher) was performed in PBS, followed by antibody staining for 30 min at 4°C in CellCover (Anacyte Laboratories) (see **Supplementary Table 3** for antibody panel and dilutions). Stained cells were sorted into PBS + 5% BSA with a FACSAria™ Fusion (BD Biosciences) equipped with a 130 μm nozzle into BSA-coated LoBind 1.5 ml tubes (Eppendorf). scRNA sequencing

#### 10x Genomics (droplet-based)

10x Genomics was performed using the Chromium Next GEM (Gel Bead-In Emulsions) Single Cell kit (v3.1, 10x Genomics) according to the manufacturer’s instructions. We pooled the T cells (CD45^high^CD34^-^CD14^-^CD235a^-^CD71^-^CD3^+^) and HSPC (CD45^dim^CD34^+^CD14^-^CD235a^-^CD71^-^) fractions in 1 lane and the stromal cells (CD34^-^ CD45^-^CD14^-^CD235a^-^CD71^-^CD38^-^) in a separate lane. Sorting and single cell processing was performed in 3 batches on different days, with each batch consisting of a mix of control, CHIP, and MDS donors to correct for batch effects. Libraries were prepared based on manufacturer’s instructions, with 13-14 PCR cycles for the final library generation. Following, the libraries were equimolarly pooled and sequenced using Illumina NextSeq 2000 with P2 and P3 flow cells.

#### CEL-Seq2 (plate-based)

The CEL-Seq2 protocol was carried out as according to published information ^169^: CD45^-^ CD14^-^CD235a^-^CD71^-^CD38^-^CD271^+^ single cells were sorted in 5 384-well plates containing 5 μl of CEL-Seq2 primer solution in mineral oil (24 bp polyT stretch, a 4 bp random molecular barcode (UMI), a cell-specific barcode, the 5*’*-Illumina TruSeq small RNA kit adapter, and a T7 promoter), provided by Single Cell Discoveries (Utrecht, the Netherlands). After sorting, the plates were centrifuged for 1 min at 1000g and immediately placed on ice and stored at −70°C. Single Cell Discoveries further processed the plates. In brief, ERCC Spike-in RNA Mix (Ambion, 0.02 μl of 1:50,000 dilution) was added to each well before cell lysis with heat shocking. Reverse transcription and second-strand synthesis reagents were dispensed using the Nanodrop II (GC biotech). After generation of cDNA from the original mRNA, all cells from one plate were pooled and the pooled sample was amplified linearly with in vitro transcription ^170^. To generate sequencing libraries, RPI-series index primers were used for library PCR. Libraries were sequenced on an Illumina Nextseq500 using 75 bp paired-end sequencing.

### SNP analysis for genotyping of donors in scRNA-seq

#### Experimental

gDNA was isolated from donors BM cells using the Macherey-Nagel NucleoSpin Tissue kit (Fisher Scientific) and was provided in a concentration ranging from 50 - 140 ng/ul for genotyping using the Infinium CoreExome-24 v1.4 BeadChip (Illumina), including 567,218 markers.

#### Computational

Raw intensity data was genotyped using the Illumina Array Analysis Platform Genotyping Command Line Interface v1.1. Resulting GTC files were converted to VCF using Illumina’s *GTCtoVCF* (v1.2.1) software (https://github.com/Illumina/GTCtoVCF) with GRCh37 reference genome and ‘--skip-indels‘ option. All VCF files were merged and indexed using *bcftools* ^171^.

To increase the number of usable genotypes, we performed imputation using the *Michigan Imputation Server* ^172^. VCF files were split by chromosome and for each of the autosomes imputation was performed using *HRC* (Version r1.1 2016) reference panel ^173^, *Eagle v2*.*4* phasing ^174^, and an *Rsq* filter threshold of 0.3. Imputed genotypes for individual chromosomes were merged and “lifted over” to GRCh38 using the LiftoverVcf program from *Picard tools* by Broad Institute (https://broadinstitute.github.io/picard/).

### scRNA-seq preprocessing and quality control

#### 10x Genomics data

Reads were aligned to the GRCh38 (v.2020-A) reference genome and quantified using *cellranger count* (10x Genomics, v.7.0.0). To assign individual cells to the corresponding donors, we used *Souporcell* ^175^ with a list of common variants from 1000 genomes project ^176^ filtered to include variants with => 2% allele frequency and a list of known genotypes derived from the SNP analysis. To increase the power of cell assignments, BAM files from individual libraries that share the same donors were combined and provided as input to *Souporcell*. Counts from *cellranger count* were adjusted for ambient RNA contamination using *SoupX* ^177^. Corrected counts were analyzed using *scDblFinder* ^178^ and used for downstream analysis using *Seurat v4*.*1*.*1* ^179^.

Cells with less than 200 genes and more than 50% mitochondrial genes per cell as well as the genes found in fewer than 3 cells were excluded from the downstream analysis. After the first round of preprocessing, normalization and clustering, we identified 2 clusters that were enriched in doublets identified by *Souporcell* **(Fig. S3E-F)**. Additionally, we identified several clusters of hematopoietic cells deriving from libraries of pooled stromal cells, namely erythroid progenitor cells, monocyte progenitor cells, and plasma cells **(Fig. S3I)**. All cells were classified as doublets by either *Souporcell* or *scDblFinder*, from doublet-enriched clusters, and clusters corresponding to hematopoietic cells deriving from the stromal libraries pools were removed and normalization, clustering and dimensionality reduction was performed again.

#### CEL-Seq2 data

CEL-Seq2 data was processed using a custom *Snakemake* ^180^ pipeline. In short, a hybrid genome reference and annotation was prepared by combining human genome reference GRCh38 and a gene annotation from Gencode Release 32 with the reference sequences and annotation of the Ambion ERCC RNA Spike-In Mix (Thermo Fisher Scientific). Raw Fastq files were aligned against the hybrid reference using *STAR aligner* (v2.7.10b) ^181^. Resulting BAM files were filtered to remove reads without an assigned cell barcode or UMI using *SAMtools* ^171^ and *GNU grep*. Filtered BAM files were used to create the gene count tables using the *dropEst* pipeline (v0.8.6) ^182^ with adjusted settings for CEL-Seq2 experiment and counting of any UMIs that overlap either exonic or intronic regions. The results were additionally corrected for UMI collisions and sequencing errors using *dropEst*. Corrected counts were used for downstream analysis and integrations using *Seurat v4*.*1*.*1* ^179^. Cells with fewer than 1000 UMI counts or belonging to a sample with fewer than 10 cells recovered were removed.

### Normalization, dimensionality reduction, and clustering scRNA-seq data

Following quality control and prior to dimensionality reduction, the raw counts were normalized using *SCTransform* v2 ^183^. 3000 most variable features were selected using SCTransform. Ribosomal, mitochondrial, sex chromosome genes and transcripts were excluded from the variable features. Selected features were used for principal component analysis (PCA) ^184^ and then uniform manifold approximation and projection (UMAP) on the first 50 principal components ^185^. Cells were then grouped into clusters using the Leiden algorithm ^185^ and the optimal clustering resolution was determined using *Clustree* ^186^.

Full dataset was classified into 3 populations of interest: HSPCs, T cells, and stromal cells; and each subset was normalized separately using SCTransfrom as described above. For integration within the HSPC and T cell populations, Seurat objects were split based on the donor ID, renormalized using SCTransform as described above, and integrated using anchor-based workflow from Seurat. Stromal population from 10x was integrated with the CEL-Seq2 data based on the experiment type using anchors integration.

Marker genes of the unsupervised clusters were identified using Seurat’s *FindAllMarkers* function. Genes considered were detected in at least 25% of cells per cluster (min.pct=0.25) and had at least 0.25 log2-fold change relative to all cells outside the cluster (logfc.threshold=0.25, only.pos=T). Differentially expressed genes between clusters were identified using likelihood ratio test for logistic regression model predicting group membership (test=“LR”).

Regulon activity quantification in scRNA-seq data Transcription factor regulons were inferred using the pySCENIC method (v.0.11.1) ^187^ via a custom *Snakemake* pipeline. Raw expression data was randomly split into 10 subsets. Initial co-expression modules were constructed using the *GRNBoost2* algorithm and enriched for transcription factor motifs using the human motif databases v10 (hg38_10kbp_up_10kbp_down_full_tx_v10_clus t.genes_vs_motifs.rankings.feather and hg38_500bp_up_100bp_down_full_tx_v10_clust.genes_vs_motifs.rankings.feather) from cisTarget (https://resources.aertslab.org/cistarget/databases/) for each of the subsets with 50 repeats each. Resulting regulons were filtered to only include target genes that were identified for a given regulon in at least 5 independent runs and combined. Regulon activities were quantified using the AUCell method of pySCENIC. For the T cells populations, regulons identified in Mathioudaki et al. ^65^ were used to compute AUCell scores in our scRNA data using the AUCell R package ^66^.

### Cell type annotation in scRNA-seq data

Cell clusters from integrated populations (HSPCs, T cells, and stromal cells) were manually annotated based on the expression of marker genes and AUCell activity scores of regulons. For the unfiltered dataset used in SpliceUp predictions, we included HSPCs that came from the sorting for Stroma and performed an additional label transfer using the *BoneMarrowMap* R package ^188^ with the reference dataset from Roy et al. ^60^. Cells corresponding to erythroid progenitors were aggregated and added to the “EryPr cluster” that we defined previously.

### Differential gene expression in scRNA-seq between conditions

Using the annotated cell types within each population, we aggregated all of the raw counts per gene per donor within each cell type. Prior to testing, genes were filtered to include only the ones that had at least 2 fragments per million present in at least half of the donors in either testing condition. We used the DESeq2 R package ^189^ to test differential expression between conditions with donor’s sex as a covariate in the design formula. The resulting estimates for effect sizes were further corrected using the *ashr* method ^190^ via DESeq2’s function *lfcShrink*. Resulting log2-fold change values were used to perform Gene Set Enrichment Analysis (GSEA) ^167^ against the Hallmark gene sets ^191^ from MSigDB ^192^ via the clusterProfiler ^193^ R package. Enriched terms with adjusted p-value <= 0.05 were used for visualization, unless specified otherwise. In comparison of iMSC between CHIP and MDS we only used data from 10x, because sample size wasn’t big enough to properly account for multiple covariates.

Curated inflammatory and HSPC-support signature scores Gene signatures were manually curated based on ^87^ for the HSPC support signature and based on ^76^ and ^194^ for the inflammatory signature. To quantify the signature scores, we used *AddModuleScore* ^195^ from the Seurat package.

### Cell-cell interactions analysis of scRNA-seq data

To infer the potential cell-cell interactions in the scRNA-seq data, we used the *NICHES* method ^89^. The full integrated dataset was split based on donor identity and for each donor-specific subset the RunNICHES function from the NICHES package was run with *Omnipath* ^196^ as a database for ligand-receptor interactions. Individual datasets were merged using the *merge*.*Seurat* function of Seurat and filtered to remove cell pairs with fewer than 5 interactions inferred. Interactions specific to cell populations of interest were selected using the *Seurat’s* FindMarkers function.

For visualization of the number of interactions per cell pair in **Fig. 6H**,**I**, the inferred number of interactions was divided by the total number of possible cell pair combinations that could come from the corresponding cell types in each donor, i.e., within each donor a product of the numbers of cells belonging to corresponding cell types.

### Single-cell trajectory inference scRNA-seq data

To construct single cell trajectories, we used the *Monocle3* ^90,197,198^ v1.3.1 R package. A subset of the integrated HSPC cells was converted to a “cell_data_set” object using the as.cell_data_set function from SeuratWrappers R package v.0.3.1. DImensionality reduction was performed using Monocle 3’s function *reduce_dimension* with max_components=3, reduction_method=“UMAP” ^199^, preprocess_method=“PCA”. Following, the cells were again clustered with the Leiden algorithm ^185^ using the *cluster_cells* function with resolution=0.01. Monocle 3’s *learn_graph* function was used to construct the trajectory graph. Three main trajectories **—** myeloid, lymphoid, and erythroid **—** were interactively constructed with Monocle 3’s *order_cells* function.

### Integration of single-cell data with public datasets

Three publicly available BM scRNA-seq datasets ^47 85 76^ were integrated with the scRNA-seq dataset from this study (see table). Stromal cells were extracted from each dataset based on author-provided annotations using the subset function in Seurat (*v5*.*0*.*3*). The datasets were merged into a single list of Seurat objects, followed by standard pre-processing, including normalization, variable feature identification, and data scaling using the default values. Principal component analysis (PCA) was initially applied as dimensionality reduction. Data integration was performed using Harmony ^200^, using the donor, dataset, and sequencing technology (10x Genomics or CEL-Seq2) as integration variables (*group*.*by*.*vars*), and PCA as the reduction method. Further dimensionality reduction was achieved using Uniform Manifold Approximation and Projection (UMAP) based on the 50 Harmony components. The Seurat layers were joined using the *JoinLayers* function from Seurat and the k nearest neighbors were computed by applying the *FindNeighbors* function. Clustering of the cells was conducted with Seurat’s *FindClusters* function, using a resolution of 0.2 to identify broad cellular populations. Additionally, the inflammation signature was calculated using the AddModuleScore function from Seurat with a manually curated list of inflammatory genes (see above, **Supplementary Table 8)**.

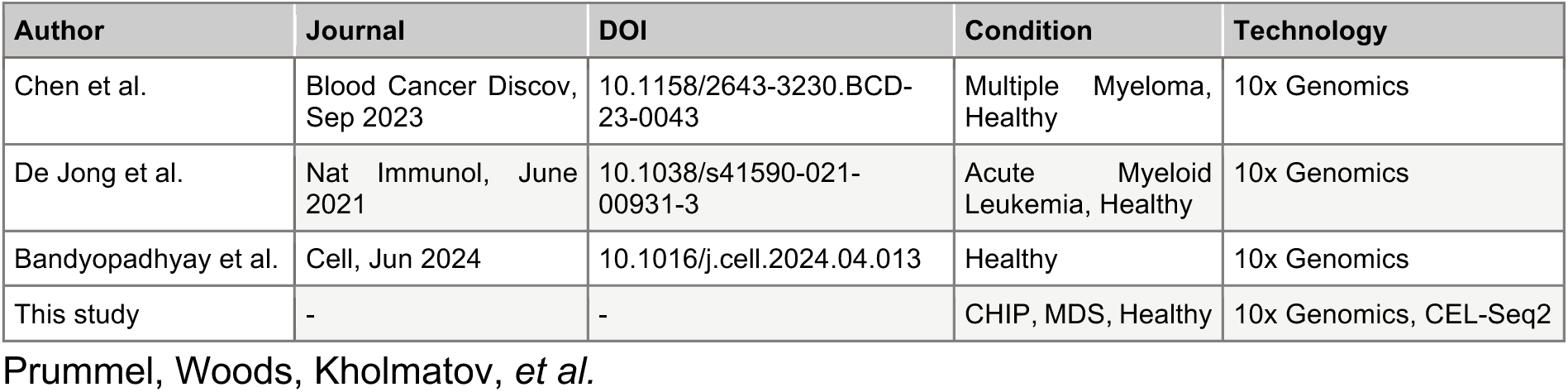

| Author | Journal | DOI | Condition | Technology |
| --- | --- | --- | --- | --- |
| Chen et al. | Blood Cancer Discov, Sep 2023 | 10.1158/2643-3230.BCD-23-0043 | Multiple Myeloma, Healthy | 10x Genomics |
| De Jong et al. | Nat Immunol, June 2021 | 10.1038/s41590-021-00931-3 | Acute Myeloid Leukemia, Healthy | 10x Genomics |
| Bandyopadhyay et al. | Cell, Jun 2024 | 10.1016/j.cell.2024.04.013 | Healthy | 10x Genomics |
| This study | - | - | CHIP, MDS, Healthy | 10x Genomics, CEL-Seq2 |

### Prediction and analysis of SF3B1-mutated cells

#### Prediction of SF3B1-mutant clones with SpliceUp

To investigate the differences between the SF3B1 wild-type (WT) and mutant clones in MDS samples, we applied the SpliceUp tool (*Kholmatov et al*., *in preparation)*. SpliceUp uses the information about known missplicing events at the 3’ splice sites associated with SF3B1 mutations in hematological malignancies to quantify the usage of normal and cryptic splice sites and use the ratio between the two to predict the mutational status of SF3B1 in individual cells. By leveraging the information across multiple gene transcripts across the genome, SpliceUp can overcome the sparsity of scRNA-seq data. In lieu of the ground truth about the mutational status of an individual cell, SpliceUp fits a multiple logistic regression model using the sample status of SF3B1 mutation as the response variable and proportions of cryptic splice site usage across multiple splice sites as the predictors. A threshold of 10% for prediction-condition fallout (proportion of false-positives among all positive predictions) was used to control the error rate of the model. To increase the power of downstream analyses, we applied SpliceUp to the subset of our 10x data that still included the erythroid progenitors from libraries sorted for stromal populations (see above under “Cell type annotation”). Additionally, we obtained preprocessed scRNA-seq data from Moura et al. ^201^ and applied SpliceUp in the same fashion.

#### Differential gene expression between SF3B1 WT and mutant clones

To quantify differential gene expression between cells predicted to be SF3B1 WT or mutant, we used Seurat’s function *FindMarkers* with test.use=“negbinom” on the subset of erythroid progenitors (**Fig. 5D**) for our data and within each of the cell types annotated in Moura et al. ^201^: HSPC, EarlyEB, MiddleEB, LateEB, Monocyte-Macrophage (**Fig. S7B**,**D**).

### Quantification and statistics

To test for significant enrichment of specific cell types in CHIP and MDS compared to controls within the scRNA-seq data set (**Fig. 3A, Fig. 5A, Fig. 6A, Supplementary Table 6**), we calculated the total number of cells for each cell type versus all other cell types within that general population (one-vs-rest) for condition of interest (CHIP or MDS) and control. Statistical significance was determined using Fisher’s exact test, followed by adjustment of P-values with the Benjamini–Hochberg procedure. Cell populations with adjusted P-values < 0.05 were highlighted as significant in the respective figure panels.

To evaluate the differences in the number of interactions between different stromal subpopulations and all HSPCs in CHIP and MDS conditions (**Fig. 4G, Fig. 6F**), we quantified the total number of interactions identified with NICHES for corresponding cell types (one-vs-rest) and condition, and performed Fisher’s exact test. P-values were corrected using the Benjamini–Hochberg procedure.

The differential HSPC-support and angiogenesis support (individual genes) of stroma subpopulations in CHIP and MDS compared to controls (**Fig. 4F, Fig. 7B**., **Supplementary Table 6**) were tested using the Wilcoxon rank-sum test.

To analyze differences in peripheral blood counts in the MDS patients across the mutational spectrum (SF3B1, other mutated splicing factors, and DNMT3A/TET2/ASXL1, **Fig. S1B, Supplementary Table 1**), we used a one-way ANOVA followed by Tukey’s HSD test for P-value adjustment. To analyze the differential abundance of the major BM populations and CD44+ MSCs (**Fig. 1, Fig. 3**., **Fig. S1, Fig. S5, Supplementary Table 4**) measured with flow cytometry across conditions, we performed a one-way ANOVA test followed by the Benjamini-Hochberg procedure for P-value correction for multiple comparisons.

For quantification of the immunofluorescence and histopathological stainings on both human and mouse bone FFPE tissue sections, multiple statistical tests were applied. We used the Kruskal-Wallis test for assessing the human BM cellularity (**Fig. 3D**), CXCL12+ MSCs (**Fig. 4C**), and vasculature quantification (**Fig. 7D**,**E**), followed up by Dunn’s test with FDR correction for multiple comparisons. One-way ANOVA with FDR correction (Benjamini-Hochberg procedure) was used for IL-1R1+ MSCs (**Fig. 3G**), the nearest distance of vasculature (**Fig. 7F**), mouse BM cellularity (**Fig. 7I**), and mouse sinusoidal and arteriolar mean vessel area (**Fig. 7L, Fig. S10F**). For the GNLY+, HSP60+, and OASL+ T cells (**Fig. 6D**) and total mouse vasculature (**Fig. 7K, Fig. S10E**), one-way ANOVA followed by Tukey’s HSD test were performed.

All statistical analyses were conducted in R or using GraphPad Prism (version 10.3.0). For ANOVA, data were tested for normal distribution (Shapiro-Wilk and Kholmogorov-Smirnov), and for sample sizes (n < 10), QQ plots were generated to guide the selection of appropriate statistical tests.

## Data and code availability

All sequencing and proteomics datasets generated for this publication will be deposited upon publication. Code used for the analyses of Nanostring nCounter gene expression data and single cell transcriptomic profiling in this paper is publicly available at https://git.embl.de/grp-zaugg/BM_CHIP_MDS. Interactive exploration of our scRNA-seq data is possible with the Shiny web application: https://shiny-portal.embl.de/shinyapps-private/app/11_cellmds. Shiny user name and password, data, and reagents are available upon request.

## Acknowledgments

We thank the tissue donors of the BoHemE study for their contribution as well as the CHOICE consortium for providing the biomaterial. We thank the tissue bank of the University Medical Clinic in Mainz for processing the samples, as well as the microscopy core facility of the Institute for Molecular Biology (IMB Mainz) for their technical support. We thank the EMBL FCCF and GeneCore facilities for technical support for the single cell experiments and Single Cell Discoveries for processing the CEL-Seq2 experiments. We thank the DKFZ Genomics and Proteomics Core Facility for the genome-wide SNP analysis. We thank the EMBL IT for providing the infrastructure and support in performing the data analysis. We thank Jean-Karim Heriche for assistance in creation and hosting of the Shiny web application. We are grateful to members from the Zaugg and Guezguez groups for their valuable input throughout the process. Additionally, we thank the DePlancke group (EPFL) for their insightful discussions. Schematics throughout the manuscript were made with BioRender (biorender.com).

## Author contributions

K.D.P., K.W., J.B.Z., and B.G. conceived the project and designed the study. K.D.P. and K.W. performed the single cell experiments. K.W. performed the co-culture and imaging experiments. M.K. processed and analyzed the scRNA-seq data, NanoString data, and integrated analysis with other datasets E.S. analyzed the NanoString data and supported experiments. E.V. integrated the publicly available scRNA-seq datasets. G.P. and K.S. performed secretomics analysis of mono- and co-cultures. M.K. and P.L.M. developed the SpliceUp algorithm. R.W. performed the NanoString experiment and M.S. supervised the original analysis. L.S.S. performed the *in vivo* genetic induction of CHIP in mice and isolated femurs. S.W. and U.O. performed diagnostic flow cytometry, retrieved clinical information, and incorporated patient data analysis. M.T. and U.P. provided administrative support and patient samples. K.D.P., M.K., J.B.Z. and B.G. analyzed and compiled the data. K.D.P., K.W., M.K., J.B.Z., and B.G. wrote the manuscript with input from all co-authors. E.L., M.S, J.J.T., J.B.Z. and B.G. supervised (certain aspects of) the project.

