## Supplementary Figures for "Inflammatory Mesenchymal Stromal Cells and IFN-responsive T cells are key mediators of human bone marrow niche remodeling in CHIP and MDS"

*[1] Molecular Systems Biology Unit, European Molecular Biology Laboratory (EMBL), Heidelberg, Germany. [2] Internal Medicine III., University Medical Center Mainz (UMC), Mainz, Germany. [3] German Cancer Consortium (DKTK), Partner Site Frankfurt/Mainz, German Cancer Research Center (DKFZ), Heidelberg, Germany. [4] Laboratory of Systems Biology and Genetics, Institute of Bioengineering, School of Life Sciences, École Polytechnique Fédérale de Lausanne (EPFL), Lausanne, Switzerland. [5] Institute of Molecular Biology (IMB), Mainz, Germany. [6] Institute of Molecular Medicine I, Proteome research, University Hospital and Medical Faculty, Heinrich Heine University Düsseldorf, Düsseldorf, Germany. [7] Molecular Proteomics Laboratory, Biological Medical Research Centre (BMFZ), Heinrich-Heine-University Düsseldorf, Düsseldorf, Germany. [8] Institute of Immunology, Faculty of Medicine Carl Gustav Carus, TU Dresden, Dresden, Germany. [9]National Center for Tumor Diseases (NCT), Partner Site Dresden, Dresden, Germany. [10] German Cancer Consortium (DKTK), Partner Site Dresden, Dresden, DKFZ, Heidelberg, Germany. [11] Department of Internal Medicine I, University Hospital Carl Gustav Carus, Faculty of Medicine Carl Gustav Carus, TU Dresden, Dresden, Germany. [12] The Jackson Laboratory, Bar Harbor, ME, USA. [13] Department of Medicine Huddinge, Center for Hematology and Regenerative Medicine, Karolinska Institute, Huddinge, Sweden. [14] University Medical Center Leipzig, Leipzig, Germany. [15] Department of Biomedicine, University Hospital Basel, University of Basel, Basel, Switzerland*

**FIGURE S1**

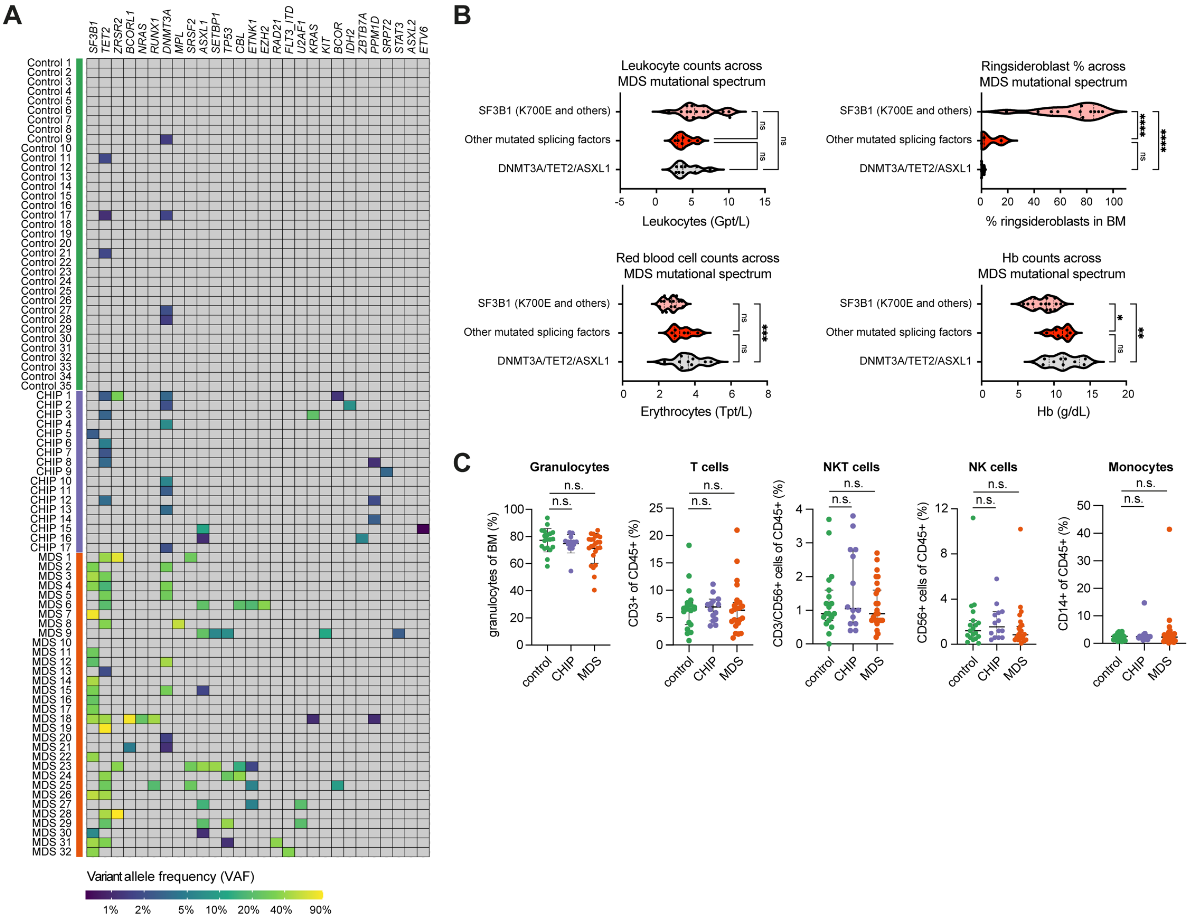


**Supplementary figure 1: Donor information and diagnostic flow cytometry of CHIP/MDS cohort** **(A)** This study included a heatmap with VAF values for mutations identified in the cohort. When several mutations are detected for the same gene within the same donor, only the maximal VAF is shown. **(B)** Violin plots representing peripheral blood counts across MDS mutational spectrum of patient's samples of the study. Median with 95% confidence intervals are shown. Statistical significance: n.s. = not significant; * q < 0.05; ** q < 0.01; *** q < 0.001, one-way ANOVA (Tukey's test correction for multiple comparisons). **(C)** Quantification of the indicated cell types' fractions in the unsorted BM (granulocytes) and CD45$^{+}$ BM (rest). Median with 95% confidence interval shown, n.s. = not significant, one-way ANOVA (FDR correction for multiple comparisons).

#### FIGURE S2

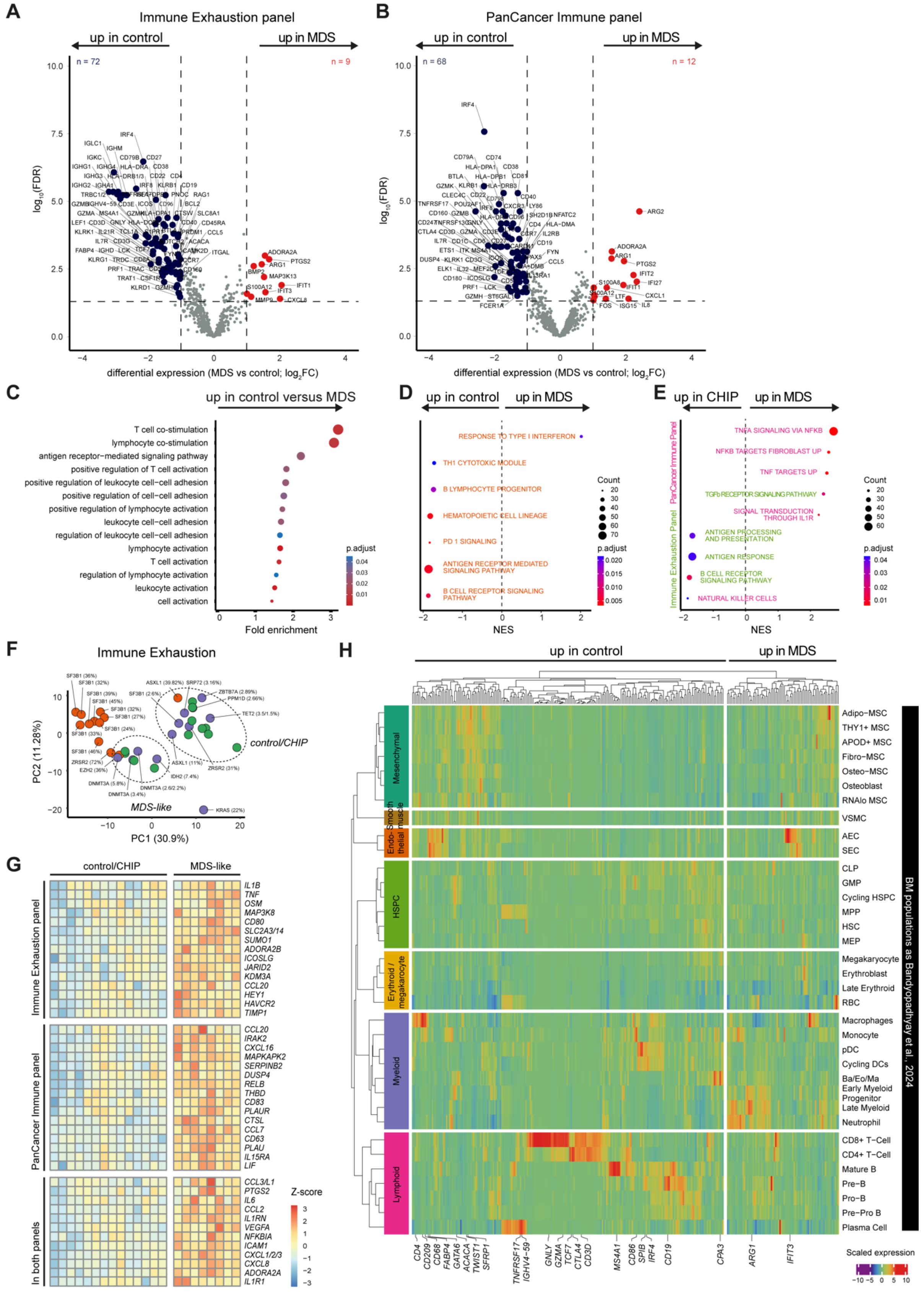


Prummel, Woods, Kholmatov *et al.*

**Supplementary figure 2: NanoString analysis of unsorted whole BM revealed hematopoietic abnormalities in MDS patients**

**(A-B)** Volcano plots of the A) Immune Exhaustion and B) PanCancer Immune Profiling panels showing A) 72 downregulated and 9 upregulated and B) 68 downregulated and 12 upregulated genes in MDS donor compared to controls (FDR 0.05). **(C)** Dot plot showing gene ontology (GO) analysis computed for all upregulated genes in control samples versus MDS samples, focused on terms specific to T cell biology and function. Statistically significant associations are highlighted by a color scheme ($p < 0.05$). **(D)** Gene set enrichment analysis (GSEA) showing the genes up- and downregulated in MDS versus control (orange terms). **(E)** GSEA for genes up- and downregulated in MDS versus CHIP across both NanoString panels, with terms for the Immune Exhaustion panel (in green) and terms for the PanCancer Immune panel (in pink). **(F)** PCA of the Immune Exhaustion panel, highlighting the variant allele frequencies (VAF) of the CHIP and MDS donors (Fig. S1A). The CHIP and control donors can be grouped into "MDS-like" and "control/CHIP" groups. **(G)** Extended heatmap of differential genes from both NanoString panels between CHIP (MDS-like) and control (control/CHIP) groups. **(H)** Heatmap displaying the expression levels of NanoString-derived genes that are differentially expressed between MDS and control samples on scRNA-seq data from healthy BM (Bandyopadhyay et al. 2024) using original cell type annotation from the authors. Genes are averaged by cell type and scaled within each cell type for visualization.

### FIGURE S3

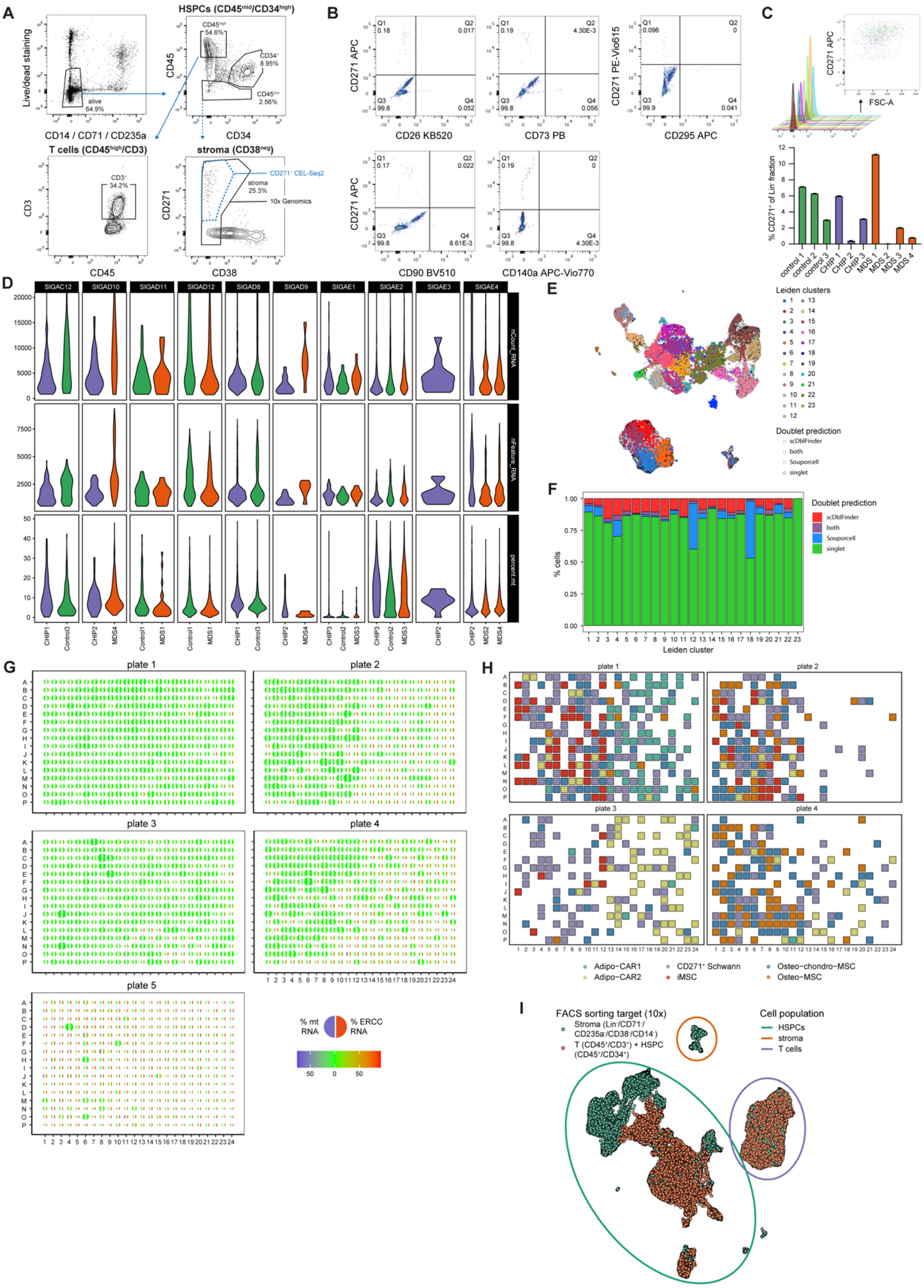


**Supplementary figure 3: FACS strategy and quality control of scRNA-seq data sets**

**(A)** Flow cytometry gating strategy for both the 10x Genomics and CEL-Seq2 scRNA-seq experiments. Schematic is showing the gating used to isolate HSPCs (CD45$^{mid}$CD34$^{+}$), T cells (CD45$^{+}$CD3$^{+}$), and stromal populations (10x Genomics: CD45$^-$CD14$^-$CD71$^-$CD235a$^-$CD38$^-$, CEL-seq2: CD271$^+$) from BM aspirates. **(B)** Surface marker analysis of stromal cells: flow cytometry dot plots displaying CD271$^+$ expression together with various markers, including CD26, CD73, CD295, CD90, and CD140a, showing CD271 labels the majority of the stromal cells compared to the other markers tested. **(C)** CD271$^+$ MSC quantification: top shows flow cytometry plot of the forward scatter (FSC-A) against CD271$^+$. Bottom shows a bar plot of counts of CD271$^+$ cells in BM aspirates from the different donors included in the scRNA-seq dataset. **(D)** Violin plots showing quantification of total counts of RNA (nCount_RNA = UMIs), number of expressed genes (nFeature_RNA = unique genes), and percentage of mitochondrial RNA (percent.mt) per cell across the different scRNA-seq libraries generated and across different donors. **(E)** UMAP visualization of Leiden clustering of scRNA-seq data, highlighting the predicted singlets and doublets in the dataset. Cells are color-coded based on the identified clusters and the doublet detection methods used (scDbFinder and Souporcell). **(F)** Bar plot displaying the proportion of the predicted singlets and doublets across the Leiden clusters in E). Two of the clusters (12 and 18) that show a very high proportion of doublets were removed from the analysis. **(G)** Dot plots showing the proportions of mitochondrial RNA (% mt RNA, blue) and ERCC RNA spike-in (% ERCC, red) for each well in the CEL-Seq2 plates (1–5). % mt RNA and % ERCC are measures for the RNA quality across wells. **(H)** Visualization of the distribution of stromal cell subtypes across the CEL-Seq2 plates that passed the quality controls (1–4), highlighting variability in cell type composition between samples. **(I)** UMAP representation of the unfiltered scRNA-seq dataset, visualizing the sorting strategy used to produce 10x libraries. HSPC contamination within libraries enriched for Stroma most notably is due to erythroid and early monocyte progenitors coming from MDS donors.

**FIGURE S4**

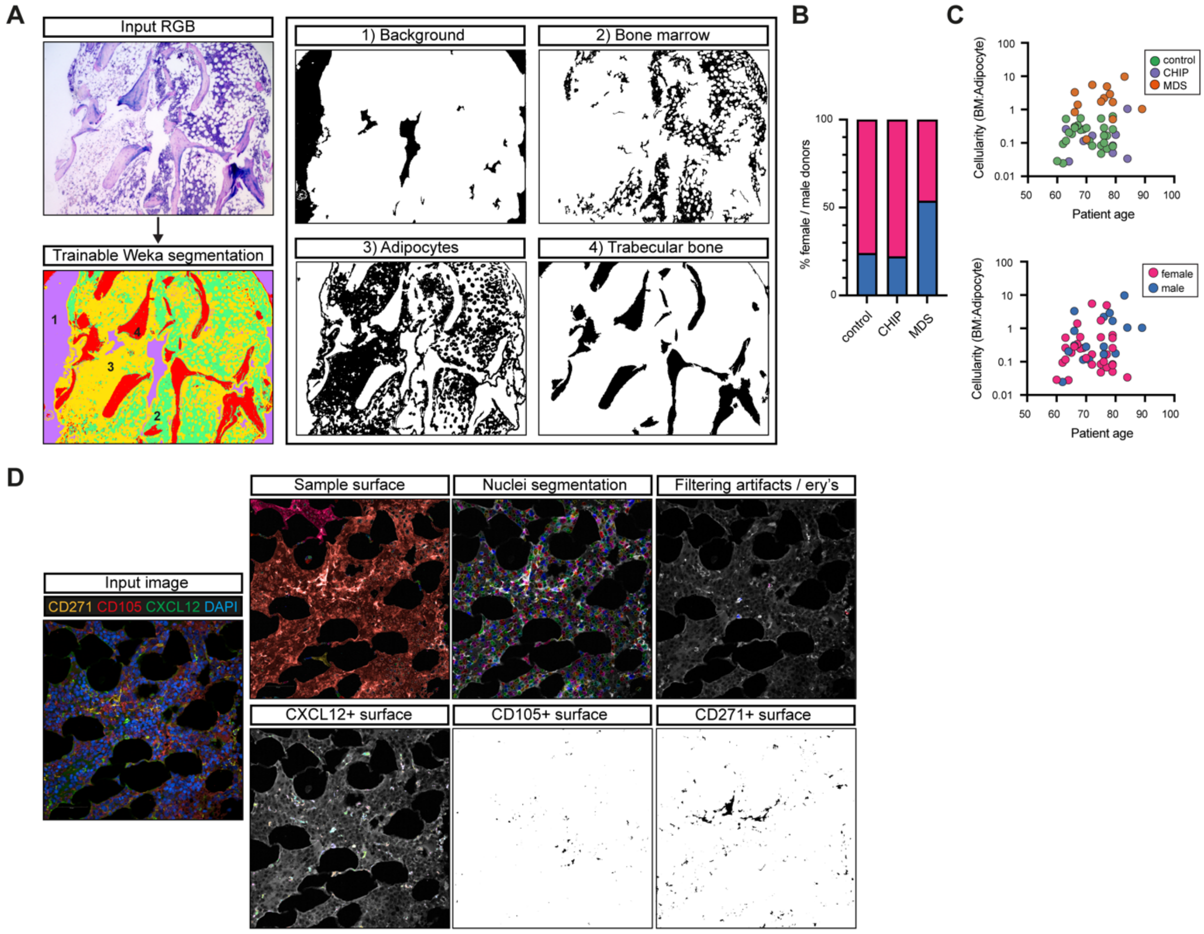


**Supplementary figure 4: Image segmentation and quantification strategy of Giemsa and immunofluorescence**

**(A)** Image sequence analysis of cellularity of Giemsa stainings. Trainable Weka-segmentation in Fiji was used to classify RGB images from stained BM. Four classes were trained: 1) background (purple), 2) bone marrow (green), 3) adipocytes (yellow), and 4) trabecular bone (red). Filtering and cleanup of the detected classes resulted in masks used for cellularity analysis in Fig. 3C (surface area bone marrow/surface area adipocytes). **(B)** Proportion of male/female donors across the cohort for the Giemsa staining and cellularity quantification. **(C)** Scatter plots visualizing the trend between cellularity, age, condition, and sex. **(D)** Semi-quantitative analysis of BM FFPE trephine samples using PerkinElmer Harmony software. For each sample, initial quality control was performed to identify fields with artifacts and uneven staining. For all remaining fields (minimum 50-300 per sample), a pipeline was set up allowing automated image analysis. Automatic thresholds were used to identify positive (surface) staining. For nuclei segmentation, in some samples, a gaussian blur on DAPI staining was employed to prevent over-segmentation. An additional check was performed to remove small artifacts and background signals from autofluorescence due to erythrocytes (ery's). With all populations of interest, overlap (%) and neighbor distance were calculated.

#### FIGURE S5

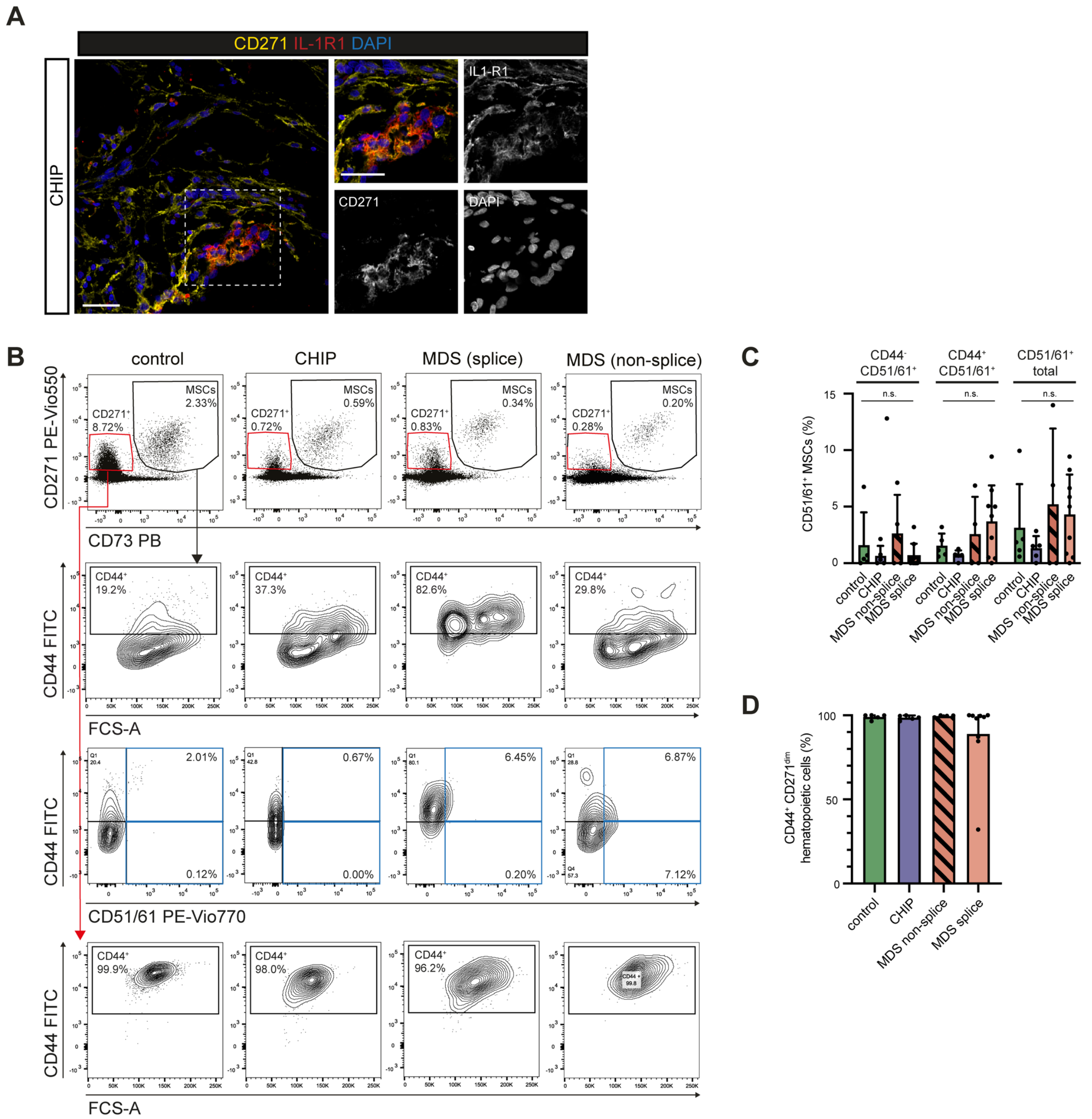


**Supplementary figure 5: Characterization strategy of iMSC by flow cytometry and tissue imaging** **(A)** Immunofluorescence images showing co-staining of IL-1R1 (red) and CD271 (yellow) in BM FFPE tissue sections from a CHIP donor. Insets highlight areas containing inflammatory IL-1R1$^{+}$CD271$^{+}$ stromal cells. DAPI (blue) stains the nuclei. Scale bars are 25 µm. **(B)** Representative dot plots visualizing the CD271$^{+}$/CD73$^{+}$ stromal population (top row, used for the analysis in Fig. 3H,I), CD44$^{+}$ MSCs (2nd row), and CD44$^{+}$/CD51/61$^{+}$ double-positive MSCs (3rd row) across control, CHIP, and MDS splice and non-splice samples as measured by flow cytometry. The last row visualizes the CD44 expression in the CD271$^{dim}$ population (highlighted in red) with low FCS-A. **(C)** Bar graph quantifying the percentage of CD44$^{+}$ and CD51/61$^{+}$ MSCs in control, CHIP, and MDS samples from B). Mean with SD shown, n.s. = not significant (two-way ANOVA, FDR correction for multiple comparisons). **(D)** Bar graph quantifying the percentage of CD44$^{+}$CD271$^{dim}$ cells.

#### FIGURE S6

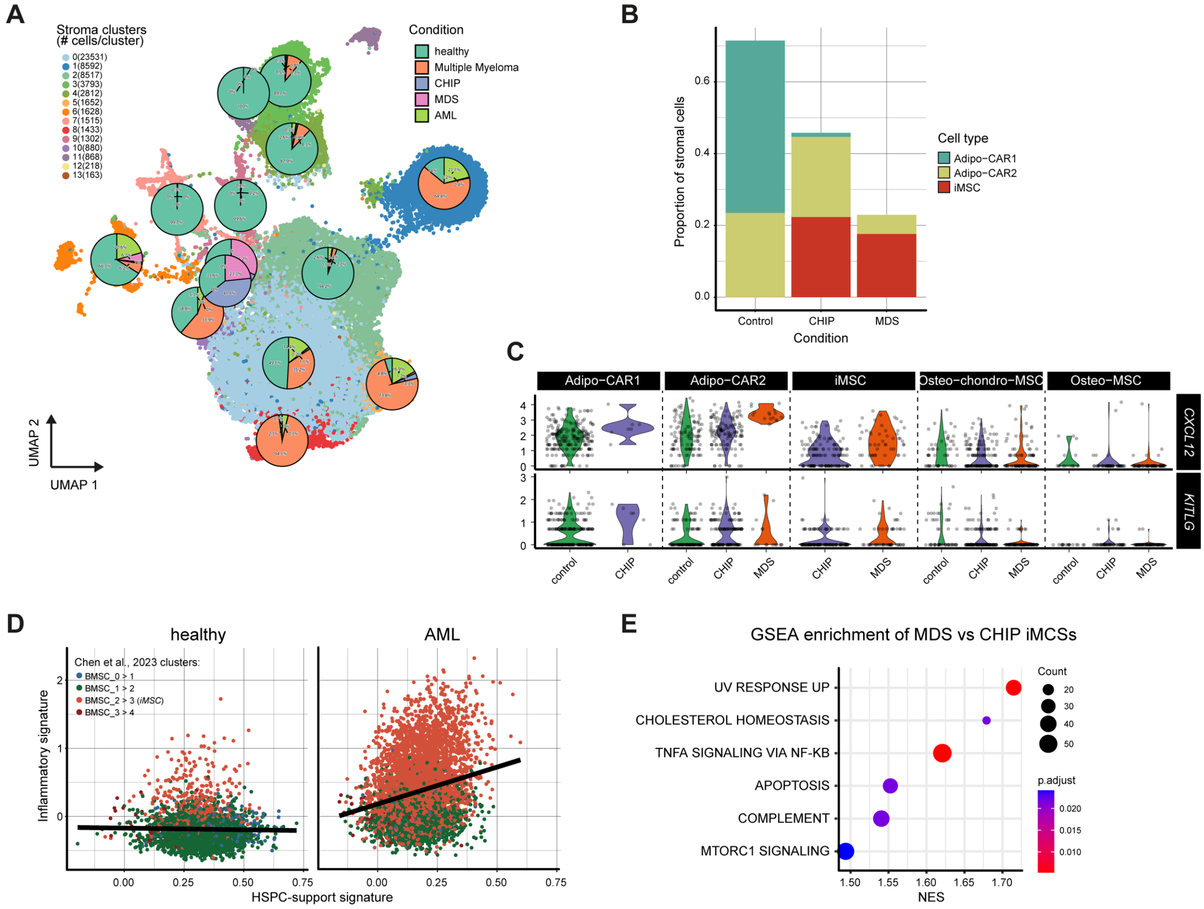

**Supplementary figure 6: Inflammatory signatures in stromal cell clusters across control, CHIP, MDS, and AML**

**(A)** Clustering of the integrated UMAP of stromal cell populations from healthy donors compiled for all 4 integrated datasets (see Fig. 3), visualizing 14 distinct stromal clusters. Pie charts indicate the contribution of different conditions (healthy, MM, CHIP, MDS, AML) to each cluster. **(B)** Bar plot visualizing the proportion of Adipo-CAR1/2 and iMSCs among all stromal cells across the different conditions (control, CHIP, MDS). **(C)** Normalized expression of *CXCL12* and *KITLG* across all stromal populations in the different conditions. **(D)** Scatter plots showing the relationship between our curated HSPC-support signature and inflammation signature scores in stromal cells of healthy and AML BM (Chen et al. 2023). Different colors represent the distinct stromal cell populations as indicated by the original publication. Black trend lines indicate the direction of the correlation in each condition. **(E)** GSEA against the Hallmarks gene sets from MSigDB, comparing the differential gene expression between iMSCs from CHIP and MDS samples.

#### FIGURE S7

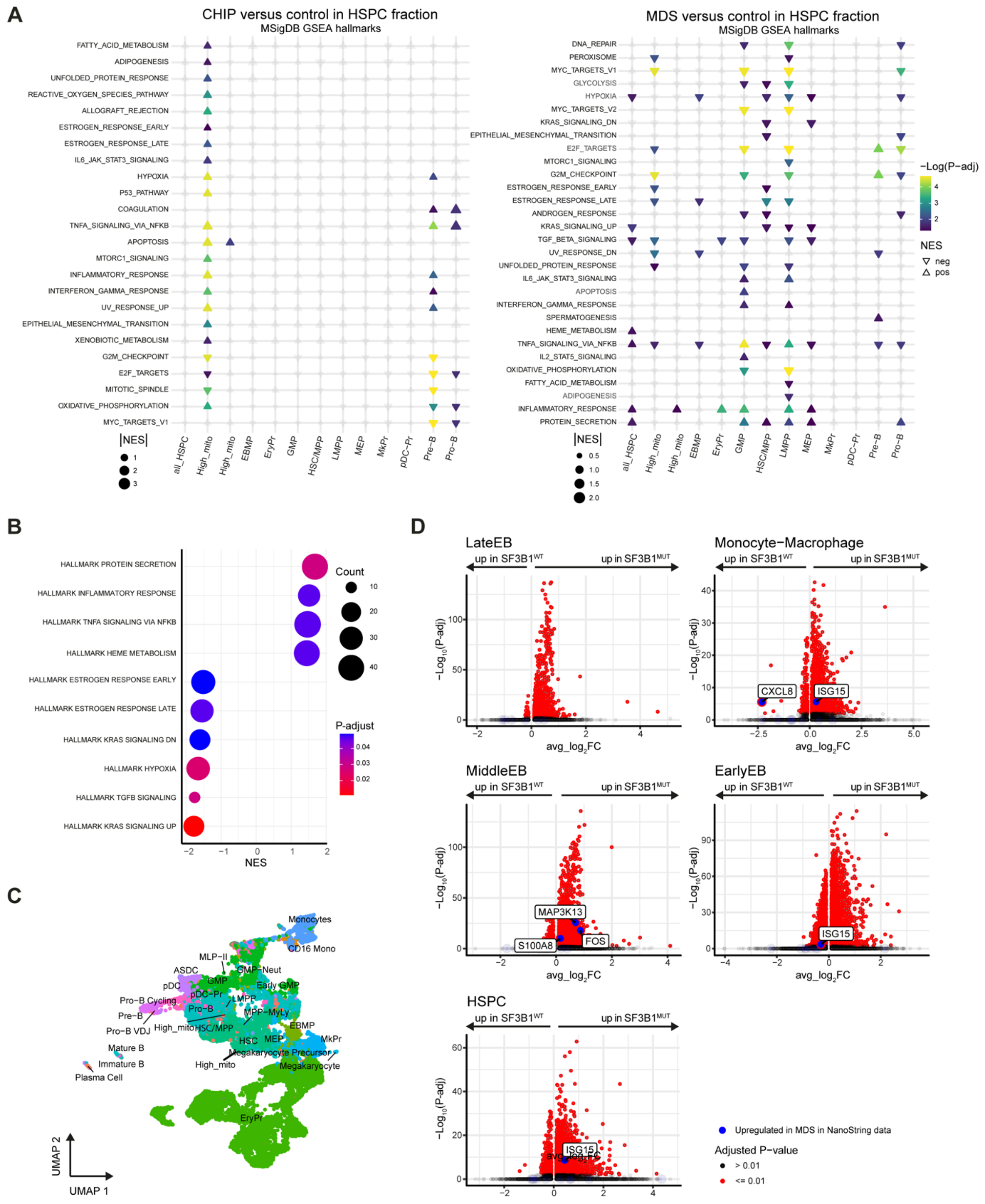

**Supplementary figure 7: Gene expression differences in HSPCs across control, CHIP, and SF3B1$^{WT}$/SF3B1$^{MUT}$ MDS**

**(A)** GSEA against the Hallmark gene sets from MSigDB, comparing differential gene expression between CHIP (left) or MDS (right) and control samples across HSPC cell types (Methods). **(B)** GSEA results for all HSPC populations (without monocytes), comparing MDS and control samples. Color scale indicates the adjusted P-value. **(C)** UMAP of the HSPC cell fraction before excluding cells from libraries enriched for stromal cells. Manually annotated cell types were combined with labels transferred from BoneMarrowMap (Roy et al. 2021). All populations corresponding to various stages of erythropoiesis were merged to perform differential expression analysis between SF3B1$^{MUT}$ and SF3B1$^{WT}$ predictions in MDS samples using SpliceUp (Fig. 5E). **(D)** Differentially expressed genes (DEG) between SpliceUp-predicted $SF3B1^{WT}$ and $SF3B1^{MUT}$ across multiple cell populations in the scRNA-seq from Moura et al.(Moura et al. 2024). DEG identified in NanoString analysis between control and MDS samples are highlighted. Statistical significance: $P < 0.01$, negative binomial test performed on single cells (Methods).

#### FIGURE S8

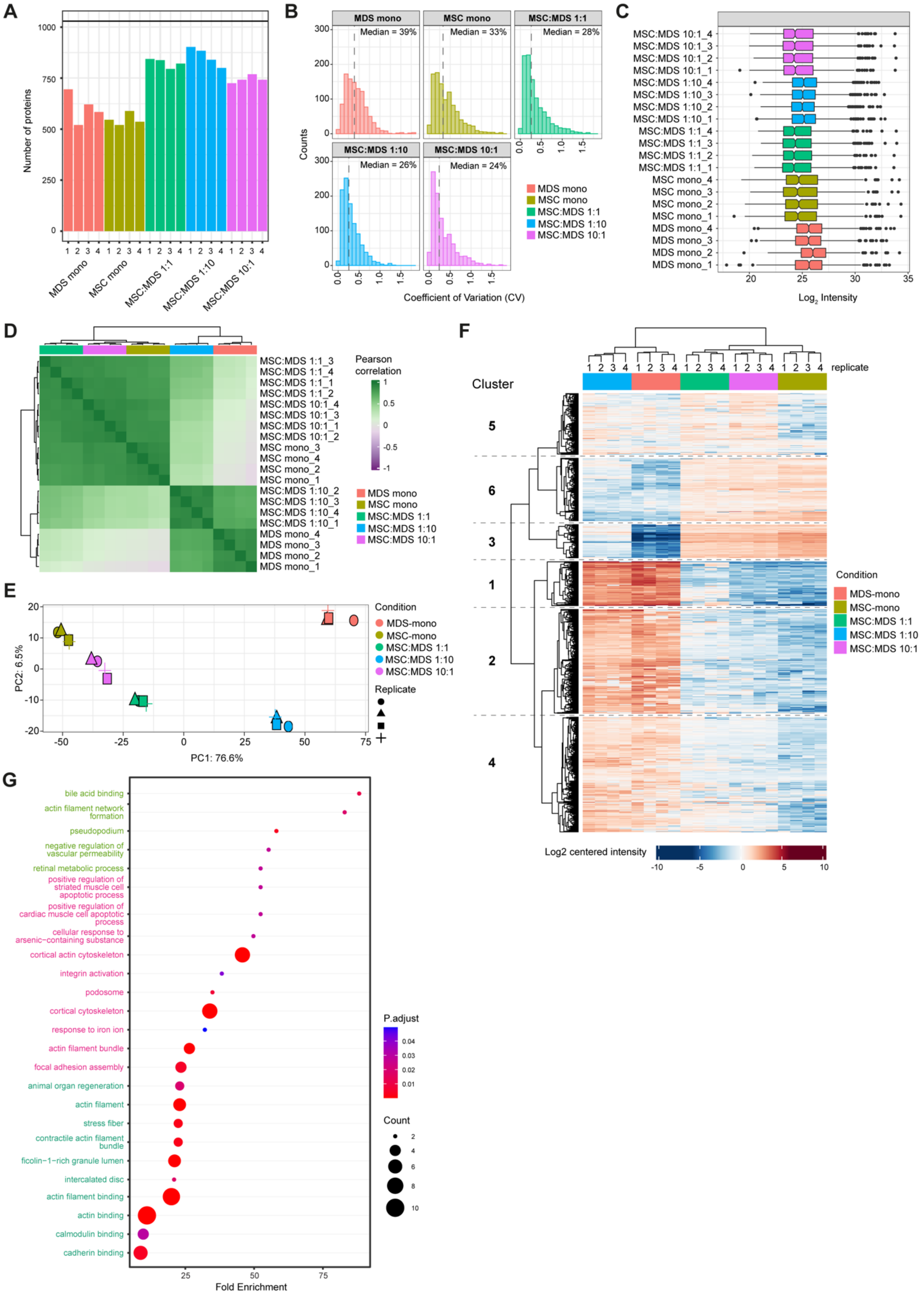


**Supplementary figure 8: Proteomics quality control**

**(A)** Number of proteins recovered per replicate (n = 4) across all conditions. **(B)** Coefficient of variation (CV) values distribution calculated for each protein for each replicate, per condition. **(C)** Data normalization based on MaxLFQ algorithm. **(D)** Correlation matrix between the replicates of the 5 different conditions using the Pearson correlation coefficient. Only proteins detected in all replicates across all conditions were included. **(E)** Principal Component Analysis (PCA) illustrating sample groups and replicates. **(F)** Heatmap showing differentially expressed proteins (rows) across all samples (columns). All pairwise comparisons (FDR and log2 fold change) have been grouped into 6 clusters based on their difference and close protein expression across all conditions. **(G)** GO term enrichment analysis of the proteins of cluster 5 in panel F).

**FIGURE S9**

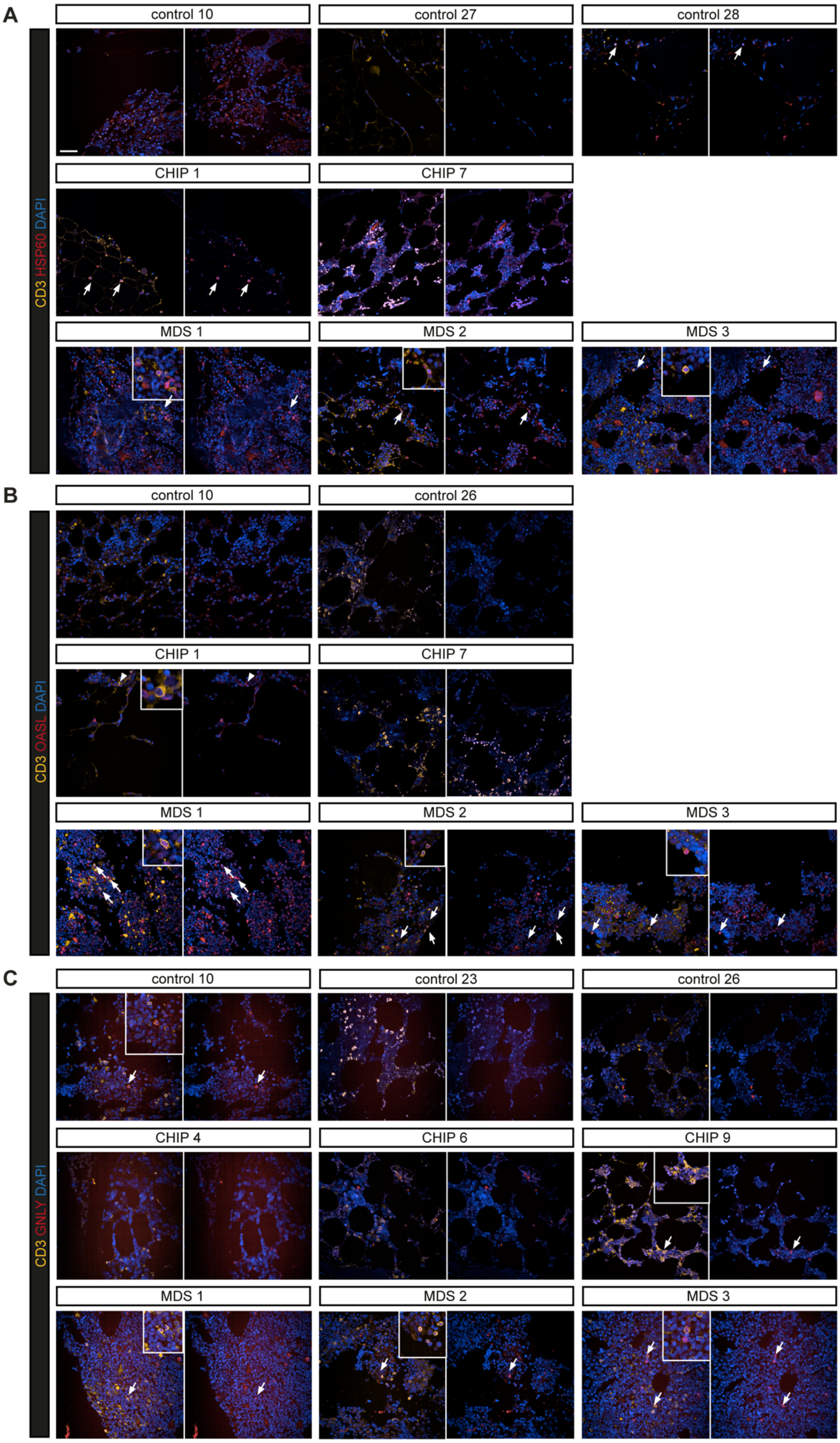


Prummel, Woods, Kholmatov *et al.*

**Supplementary figure 9: Visualization of HSP+, IFN response, and $T_{EMRA}$ cytotoxic T cells in the BM of CHIP and MDS**

**(A-C)** Representative images of CD3 T cells (yellow), co-stained with the cell cluster markers HSP60 (red, A), OASL (red, B), or GNLY (red, C). Zoomed-in regions highlight and arrows indicate double-positive cells. Scale bar is 25 µm.

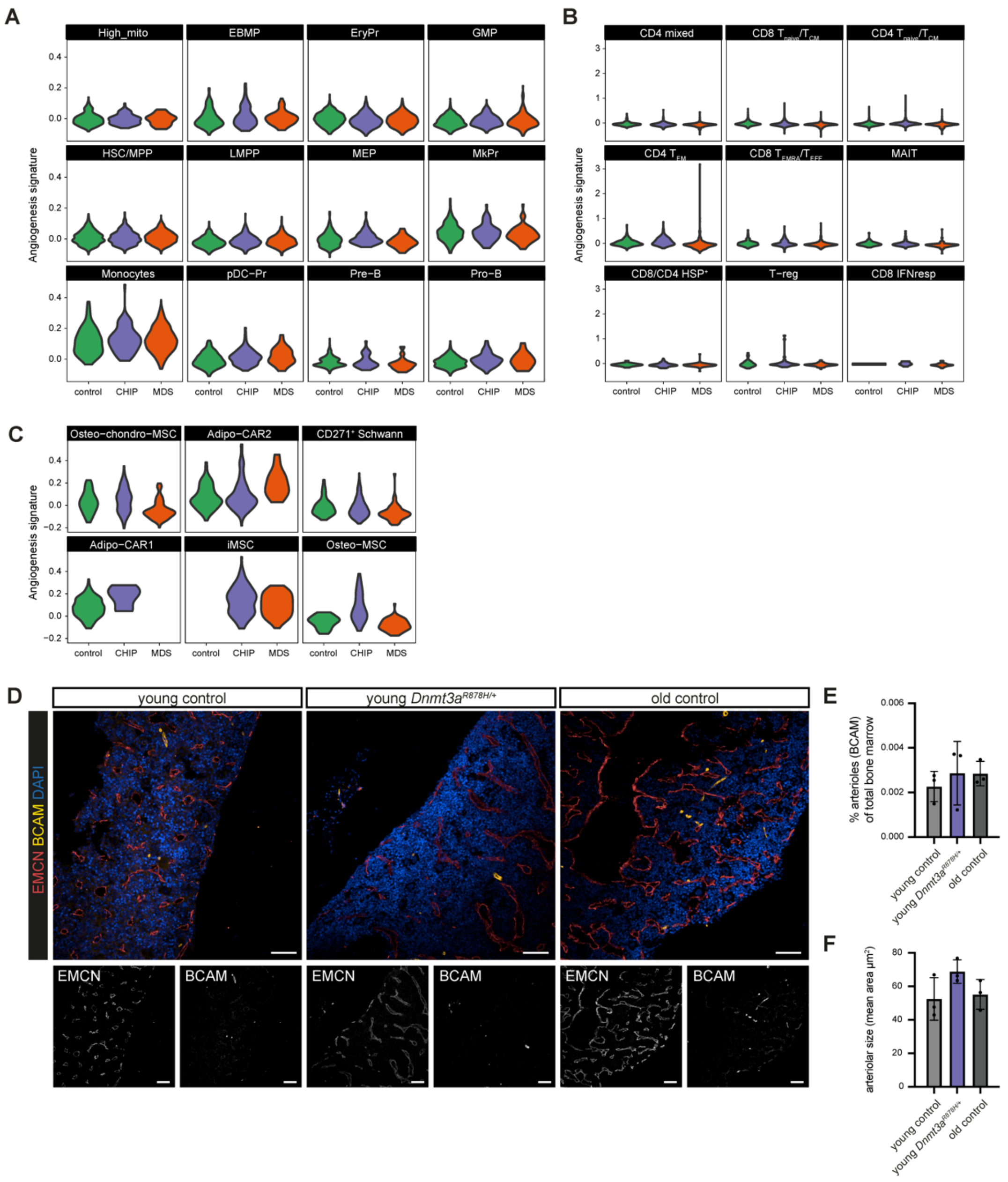


**Supplementary figure 10: Angiogenesis scores across all BM stromal, HSPCs, and T cell clusters and vascular remodeling characterization in CHIP**

**(A-C)** Volcano plots representing the angiogenesis signatures in all individual A) HSPC clusters, B) T cell clusters, and C) stromal clusters across control, CHIP, and MDS samples from the scRNA-seq data. **(D)** Representative images of mouse FFPE bone tissue section co-stained with sinusoidal (EMCN) and arteriolar (BCAM) markers of young control, young *Dnmt3a*$^{R878H}$, and old control mice. DAPI (blue) stains the nuclei. Scale bar is 100 µm. **(E)** Quantification of the total arteries/arteriolar (BCAM) area in the mouse femur. Statistical significance: ns, adjusted P-value < 0.05, two-way ANOVA (Tukey's test for multiple comparison correction). **(F)** Quantification of the mean area of the arteries/arteriolar vasculature (arteries/arteriolar size). Statistical significance: ns, adjusted P-value < 0.05, one-way ANOVA (FDR for multiple comparison correction).
